# FMRP promotes RNA localization to neuronal projections through interactions between its RGG domain and G-quadruplex RNA sequences

**DOI:** 10.1101/784728

**Authors:** Raeann Goering, Laura I. Hudish, Bryan B. Guzman, Nisha Raj, Gary J. Bassell, Holger A. Russ, Daniel Dominguez, J. Matthew Taliaferro

## Abstract

The sorting of RNA molecules to distinct subcellular locations facilitates the activity of spatially restricted processes through local protein synthesis. This process affects thousands of transcripts yet precisely how these RNAs are trafficked to their destinations remains generally unclear. Here we have analyzed subcellular transcriptomes of FMRP-null mouse neuronal cells to identify transcripts that depend on FMRP for efficient transport to neurites. We found that these FMRP RNA localization targets contain a large enrichment of G-quadruplex sequences, particularly in their 3′ UTRs, suggesting that FMRP recognizes these sequences to promote the localization of transcripts that contain them. Fractionation of neurons derived from human Fragile X Syndrome patients revealed a high degree of conservation in the identity of FMRP localization targets between human and mouse as well as an enrichment of G-quadruplex sequences in human FMRP RNA localization targets. Using high-throughput RNA/protein interaction assays and single-molecule RNA FISH, we identified the RGG domain of FMRP as important for both interaction with G-quadruplex RNA sequences and the neuronal transport of G-quadruplex-containing transcripts. Finally, we used ribosome footprinting to identify translational regulatory targets of FMRP. The translational regulatory targets were not enriched for G-quadruplex sequences and were largely distinct from the RNA localization targets of FMRP, indicating that the two functions can be biochemically separated and are mediated through different target recognition mechanisms. These results establish a molecular mechanism underlying FMRP-mediated neuronal RNA localization and provide a framework for the elucidation of similar mechanisms governed by other RNA-binding proteins.

## INTRODUCTION

Essentially all eukaryotic cells contain subcellular regions that are associated with specific functions. The identity and activity of these regions are in large part defined by the proteins contained within them. Cells require an efficient sorting process to get proteins required for function to their correct location. For many genes, this fundamental problem is solved by transporting the RNA molecule encoding the protein, rather than the protein itself, to the desired location. The transcript is then translated on-site to immediately produce a correctly localized protein.

RNA localization has been studied extensively in diverse systems (Ephrussi et al., 1991; Long et al., 1997; Moor et al., 2017). Thousands of transcripts are asymmetrically distributed within cells and therefore subject to localization regulation (Cajigas et al., 2012; Gumy et al., 2011; Lécuyer et al., 2007; Taliaferro et al., 2016a; Taylor et al., 2009). However, precise molecular mechanisms that govern how transcripts are transported have been worked out in detail for only a small subset of heavily studied localized RNAs (Bertrand et al., 1998; Ephrussi et al., 1991; Jambhekar and Derisi, 2007; Kislauskis et al., 1994). In these RNAs, sequences that mark transcripts for transport, sometimes termed “zipcodes”, are often found in 3′ UTRs (Jambhekar and Derisi, 2007). Zipcode sequences are then bound by RNA-binding proteins (RBPs) that mediate RNA transport (Farina et al., 2003; Huang et al., 2003; St Johnston et al., 1991; Zhang et al., 2001). The zipcode/RBP pairs that are known were often identified through careful but laborious reporter experiments that involve systematic deletions or mutations that inactivate putative regulatory elements.

A large disparity persists, though, between the number of RNAs that are targets of localization regulation and the number of known localization regulatory elements within those RNAs. High-throughput experiments that probe the localization of thousands of transcripts at once may be able to help bridge this gap. By providing many examples of transcripts whose localization is regulated by a given RBP, commonalities among those transcripts can shed light on general rules that define sequence features that regulate RNA localization as well as the RBP that binds them.

FMRP is an RBP that is highly expressed in the brain and, and loss of FMRP expression has long been known to be the cause of Fragile X syndrome (FXS) (Santoro et al., 2012). Loss of expression of the FMR1 gene in humans is associated intellectual disabilities and occurs in approximately 1 in 5000 males (Coffee et al., 2009). FMRP-null mice display similar phenotypes (Kazdoba et al., 2014). FMRP has been shown to regulate RNA metabolism at the level of translational repression and RNA localization (Darnell et al., 2011; Dictenberg et al., 2008). The relative contribution of these activities to observed phenotypes is generally unclear. Although genome-wide studies probing the translation repression activity of FMRP have been performed (Darnell et al., 2011), much less is known about the RNAs that depend on FMRP for efficient localization to neuronal projections. FMRP has been implicated in the transport of a handful of RNAs (Dictenberg et al., 2008; Muddashetty et al., 2007; Zalfa et al., 2007), but there is no general understanding of the identity of transcripts whose localization is regulated by FMRP.

How FMRP recognizes and binds its functional targets is also not well understood. Several studies have reported transcript and RNA motif targets for FMRP in vitro and in vivo (Ascano et al., 2012; Brown et al., 2001; Darnell et al., 2001, 2005a; Vasilyev et al., 2015), yet considerable disagreement among these studies remains about the identity of RNA sequences that interact with FMRP with high-affinity (Anderson et al., 2016). RNA G-quadruplex sequences have been repeatedly identified as binding targets of FMRP (Darnell et al., 2001; Schaeffer et al., 2001), but other studies have found that FMRP binds completely different short RNA motifs (Ray et al., 2013) while still others seem to show that FMRP displays almost no sequence specificity at all and instead binds all along the coding sequence of its target transcripts (Darnell et al., 2011). Whether or not these potentially different modes of binding are related to the different functions of FMRP is again unknown.

## RESULTS

### Identification of transcripts that depend upon FMRP for efficient neuronal transport

We and others have previously used subcellular fractionation followed by high-throughput sequencing to quantify transcripts in the soma and neurites of neuronal cells (Taliaferro et al., 2016a; Zappulo et al., 2017). In these experiments, neuronal cells are plated on porous membranes (**Fig 1A**). The pores of these membranes are big enough to allow neurite growth through them but small enough to restrict soma growth to the top of the membrane. This allows the cells to be mechanically fractionated into soma and neurite fractions. RNA can then be isolated from both fractions and analyzed by high-throughput sequencing.

**Figure 1.**
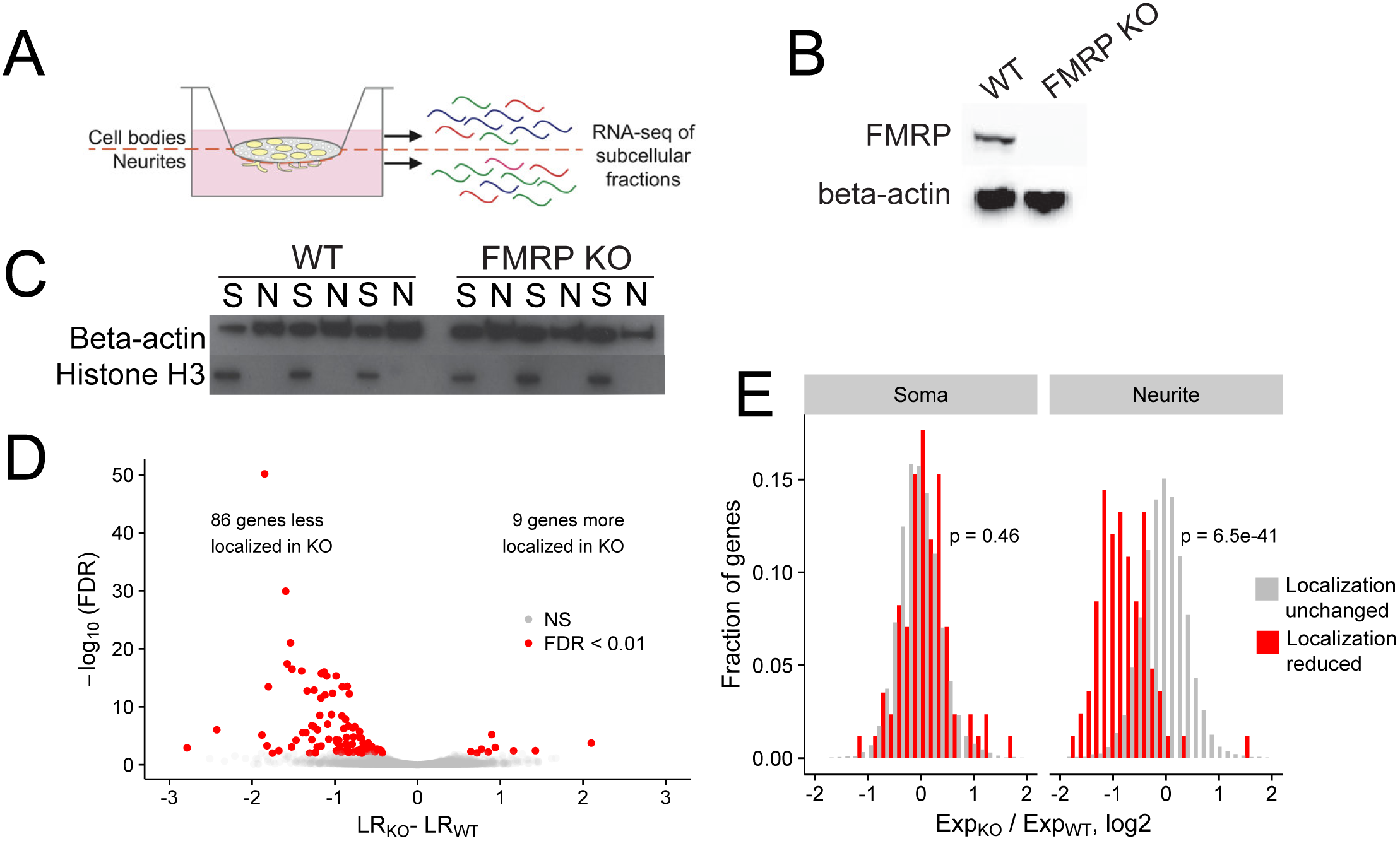
Identification of FMRP localization targets. (A) Schematic of soma/neurite fractionation. Cells are plated on porous membranes. Neurites grow down through the pores, and cells are then mechanically fractionated by scraping. (B) Western blot of wildtype and CAD FMRP knockout cells. (C) Western blot of fractionated soma (S) and neurite (N) samples. Beta-actin is a marker of both soma and neurite fractions while Histone H3, being restricted to the nucleus, is a marker of soma fractions. (D) Localization ratio (LR) comparison for all expressed genes in wildtype and FMRP-null cells. Changes in LR values and FDR values were calculated using Xtail. (E) Expression changes across genotypes in the soma and neurite fractions for FMRP localization targets (red) and nontargets (gray). P-values were calculated using a Wilcoxon rank-sum test.

Given the utility of this technique, we then set out to define transcripts whose efficient localization to neurites depended on FMRP. Using CRISPR/Cas9, we created an FMRP-null CAD mouse neuronal cell line. CAD cells are derived from a mouse brain tumor and can be induced by serum starvation to display features of neuronal differentiation, including cell cycle downregulation, expression of neuronal markers, and the appearance of neuronal morphologies (Qi et al., 1997). The CAD FMRP knockout line was found to have a single basepair deletion in all alleles of Fmr1 (**Fig S1A**) that caused a frameshift and a premature termination codon two codons downstream of the gRNA cut site. The premature termination codon was upstream of all identified RNA-binding domains in FMRP. The knockout cells displayed no detectable FMRP expression by immunoblotting, even when probed with a polyclonal FMRP antibody (**Fig 1B**).

We then fractionated wildtype and FMRP knockout CAD cells into soma and neurite fractions in triplicate. To monitor the efficiency of the fractionation, we assayed the expression of beta-actin, which should be present in all fractions, and histone H3, which, being nuclear, should be restricted to the soma (**Fig 1C**). RNA was isolated from each fractionation and profiled by high-throughput sequencing.

The expression of Fmr1 RNA was downregulated in FMRP knockout samples, as would be expected since the Cas9-induced lesion likely made the RNA a target of nonsense-mediated decay (**Fig S1B**). Principal component analysis of gene expression values showed that replicates clustered together and samples were separated along the first two principal components by compartment and genotype, respectively (**Fig S1C, D**). To quantify RNA localization, we defined a metric termed localization ratio (LR) as the log2 of a gene’s abundance in the neurite fraction divided by its abundance in the soma fraction. Positive LR values are therefore associated with neurite localization, and higher LR values are associated with stronger neurite localization.

Previous literature has demonstrated that transcripts encoding ribosomal proteins and components of the electron transport chain are strongly enriched in neurites (Gumy et al., 2011; Moccia et al., 2003; Taliaferro et al., 2016a). We therefore sought to assay the efficiency of our fractionation and RNA isolation by looking at the LR values of these genes. Reassuringly, we found that, in all samples, mRNAs encoding these proteins were strongly neurite-enriched (**Fig S1E**).

To identify genes whose transcripts depend on FMRP for efficient transport to neurites, we compared the LR values of genes between the wildtype and knockout samples using the R package Xtail (Xiao et al., 2016). Xtail is designed to analyze ribosome profiling data to identify genes whose translational efficiency changes across conditions. However, since both translational efficiency and LR are ratios of gene expression values, we reasoned that Xtail would be a good tool to identify genes whose LR value changes across samples. Using Xtail and a false discovery rate cutoff of 0.01, we identified 86 genes whose transcripts displayed impaired neurite localization, and thus decreased LR values, in the FMR1 knockout (**Fig 1D, S1F, Table S1**). We defined these genes as FMRP localization targets.

Gene ontology analysis of these targets revealed that there were enriched for genes that regulate vesicle transport in neurons (**Fig S1G**). In particular, mRNAs encoding several kinesins were less localized in the FMRP knockout. Interestingly, mRNAs encoding kinesins have been recently identified as both bound and transported by FMRP in glial cells (Pilaz et al., 2016).

Because LR is a ratio of neurite to soma gene expression values, in principle, a decrease in LR could be due to either a decrease in neurite expression or an increase in soma expression. To investigate this further, we compared the expression of the mislocalized genes in the soma and neurite compartments separately. We found that while the expression of FMRP localization targets did not change in the soma between the wildtype and FMRP knockout samples, their expression in the neurites of FMRP knockout cells was significantly reduced (**Fig 1E**). Because the soma contains the vast majority of cellular RNA, this indicates that the overall expression and processing of the localization targets is not impaired yet their ability to be trafficked to neurites is reduced. These findings suggest that these genes are in fact misregulated at the level of RNA localization.

### FMRP localization targets have significant enrichments of G-quadruplex sequences in their 3′ UTRs

After identifying functional RNA localization targets of FMRP, we then looked for commonalities among them that can give insight into how FMRP recognizes and binds them. If FMRP were directly regulating the localization of these targets, we would expect that FMRP directly binds them. To test this, we analyzed data from two previously published FMRP CLIP-seq datasets (Ascano et al., 2012; Darnell et al., 2011). Given the lack of suggesting consensus between multiple published FMRP binding datasets (Anderson et al., 2016), we defined transcripts bound by FMRP as those that appeared in both CLIP-seq datasets (Ascano et al., 2012; Darnell et al., 2011). FMRP localization targets were approximately 3-fold more likely to be directly bound by FMRP than expected by chance, that FMRP directly regulates the localization of these transcripts (**Fig 2A**).

**Figure 2.**
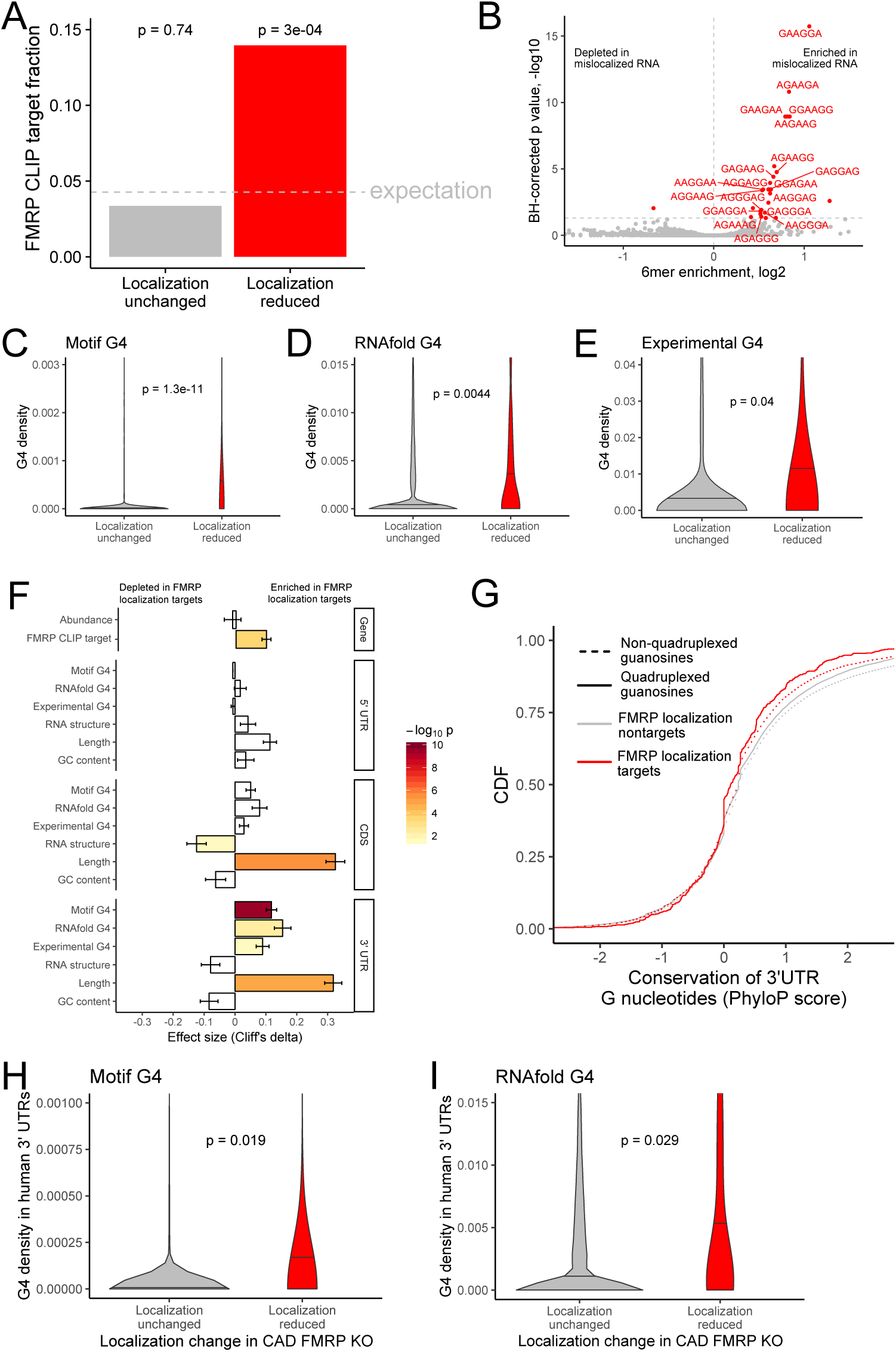
Identification of transcript features associated with the regulation of localization by FMRP. (A) Fraction of FMRP localization targets (red) and nontargets (gray) that were previously identified as directly bound by FMRP in cells. FMRP-bound RNAs were defined as the intersection between two previously published datasets (Ascano et al, 2012; Darnell et al 2011). The null, expected fraction is represented by a gray dotted line. P values were calculated using a hypergeometric test. (B) Sixmer enrichments in the 3′ UTRs of FMRP localization targets vs nontargets. Significantly enriched 6mers (BH-adjusted p < 0.05, Fisher’s exact test) are represented in red. (C-E) G-quadruplex sequence densities in the 3′ UTRs of FMRP localization targets (red) and nontargets (gray). In (C), G-quadruplex sequences were defined by a regular expression that contained for WGGA sequences separated by linkers of length 0-7 nt. Values represent sequence matches per nt. In (D), RNAfold was used to identify G-quadruplex sequences. Values represent the number of guanosine residues participating in quadruplex sequences per nt. In (E), experimentally defined G-quadruplex sequences were used. Values represent the number of basepairs that overlap with an experimentally defined G-quadruplex sequence per nt of UTR sequence. (F) Effect sizes and significance of the relationships between transcript features and whether or not a transcript’s localization was regulated by FMRP. (G) Conservation of guanosine residues in 3′ UTRs as defined by PhyloP scores. Guanosines were separated based on whether or not they were predicted by RNAfold to participate in quadruplexes and by whether or not they were contained in FMRP localization targets. (H) G-quadruplex sequence density in the 3′ UTRs of the human orthologs of FMRP targets. G-quadruplexes were defined by the regular expression used in (C), and values represent quadruplex sequences per nt. (I) G-quadruplex sequence density in the 3′ UTRs of the human orthologs of FMRP targets. G-quadruplex sequences were defined by RNAfold as in (D), and values represent the number of guanosine residues participating in quadruplex sequences per nt.

We then reasoned that sequences that were enriched in FMRP localization targets relative to nontargets could represent preferred binding sites for FMRP. Because the majority of RNA sequence elements that regulate localization identified to date reside in 3′ UTRs (Jambhekar and Derisi, 2007), we compared the abundance of all 6mers in the 3′ UTRs of FMRP localization targets and nontargets. We observed a strong enrichment for the RNA sequence ‘GGA’ in the 3′ UTR of FMRP localization targets (**Fig 2B**). GGA had previously been identified from an FMRP CLIP-seq dataset as a preferred FMRP binding motif, giving us further confidence that our identified localization targets were directly bound and regulated by FMRP (Ascano et al., 2012).

Tandem repeats of GGA sequences are known to have the ability to fold into G-quadruplex sequences (Suhl et al., 2014). G-quadruplexes are four-stranded nucleic acid structures in which guanosine residues interact with each other across the strands (Rhodes and Lipps, 2015). Given that G-quadruplex structures have been shown to be specifically bound by FMRP *in vitro*, (Didiot et al., 2008; Vasilyev et al., 2015), we then asked if G-quadruplex sequences were enriched in FMRP localization target sequences. To do so, we employed two computational methods to predict G-quadruplex sequences from RNA sequences. Using a regular expression we searched for four repeated WGGA sequences separated by 0-7 nucleotide linkers (Huppert and Balasubramanian, 2007) where W represents either adenosine or uracil. We also used the RNA secondary structure prediction software RNAfold (Lorenz et al., 2011). Both methods agreed that the 3′ UTRs of FMRP localization targets contained significantly more G-quadruplex sequences than nontargets (**Fig 2C, D**).

We then complemented these computational predictions of G-quadruplex sequences with previously published experimentally defined G-quadruplex sequences (Guo and Bartel, 2016). These were identified by quantifying reverse transcriptase stop sites in the presence of either potassium or lithium. Because potassium stabilizes G-quadruplexes and lithium destabilizes them, stop sites that display differential abundance between the two conditions are likely due to G-quadruplexes. These experimentally defined G-quadruplexes displayed many expected properties, including a large enrichment for guanosine nucleotides (**Fig S2A**), highly stable predicted secondary structures (**Fig S2B**), and a strong enrichment for predicted G-quadruplexes using both the regular expression and RNAfold methods (**Fig S2C, D**). The agreement between the computational and experimental approaches to G-quadruplex identification gave us confidence in the ability of both techniques. As with the computational predictions, we observed a significant enrichment of experimentally defined G-quadruplexes in the 3′ UTRs of FMRP localization targets (**Fig 2E**).

We then asked what other transcript features were associated with FMRP localization targets. We observed that FMRP localization targets had much longer open reading frames and 3′ UTRs than expected (**Fig 2F**), consistent with previous reports that 3′ UTR extensions and alternative 3′ UTRs can regulate RNA localization in neurons (Taliaferro et al., 2016a; Tushev et al., 2018). We observed no relationship between the overall abundance of a transcript and whether or not it was an FMRP localization target. The enrichment of G-quadruplexes in FMRP localization targets was restricted to their 3′ UTRs as we did not observe a significant relationship between CDS or 5′ UTR G-quadruplex density and localization regulation by FMRP (**Fig 2F**).

### G-quadruplexes are not positionally conserved but tend to be present in orthologous genes

The enrichment of G-quadruplex sequences in FMRP localization targets and the enrichment of those targets in FMRP CLIP-seq datasets suggested that the G-quadruplex sequences may be functional. If they were functional, we would expect them to be conserved. To test the conservation of G-quadruplex sequences, we compared the conservation of 3′ UTR guanosine residues predicted by RNAfold to participate in quadruplex interactions to those not predicted to be in quadruplexes. Surprisingly, we found that guanosines in quadruplexes were less conserved than those not in quadruplexes, and those in FMRP localization targets were less conserved than those in nontargets (**Fig 2G**). However, this analysis relies on elements being positionally conserved such that they can be aligned across genomes. Elements within UTRs do not have to be positionally conserved, only present, to be functional.

To assess this, we calculated G-quadruplex densities in the 3′ UTRs of the human orthologs of the FMRP localization targets and nontargets. We observed a significant enrichment of G-quadruplexes in the human orthologs of the FMRP localization targets, arguing that their function may be conserved across species (**Fig 2H, I**).

### Mislocalized transcripts in FXS patient-derived neurons are enriched for G-quadruplex sequences

To more directly test the localization regulatory activity of FMRP in human cells, we created induced pluripotent stem (iPS) cell lines from FXS and unaffected patient samples. We differentiated these cells over the course of 20 days into motor neurons (Chambers et al., 2009; Du et al., 2015; Reinhardt et al., 2013) (**Fig S3A**). We then plated these cells on porous membranes and performed the same subcellular fractionation and sequencing experiment that was performed on the CAD cell samples using four technical replicates.

As expected, the FXS neurons displayed a large decrease in both FMR1 RNA and FMRP protein (**Fig 3A, B**). We assayed the efficiency of the fractionation as before by immunoblotting (**Fig S3B**) and looking for the enrichment of mRNAs encoding ribosomal proteins and components of the electron transport chain in the neurite fraction (**Fig S3C**) and found that both performed as expected. PCA analysis of gene expression values showed a separation of samples by compartment and genotype along the first two principal components (**Fig S3D**). We first looked at the localization of the human orthologs of the FMRP localization targets that were identified in the mouse CAD system. Strikingly, the human orthologs of the FMRP localization targets were significantly less neurite-localized in FXS neurons than in unaffected neurons, indicating that the functional localization targets of FMRP were broadly conserved across species (mouse and human) and experimental systems (neuronal cell line and iPS-derived neurons) (**Fig 3C**). Overall, we observed a modest (Spearman R = 0.21) but highly significant (p = 1e-37) correlation between the mislocalization of genes in the CAD knockout system and the mislocalization of their human orthologs in the FXS iPS system (**Fig S3E, Table S2**).

**Figure 3.**
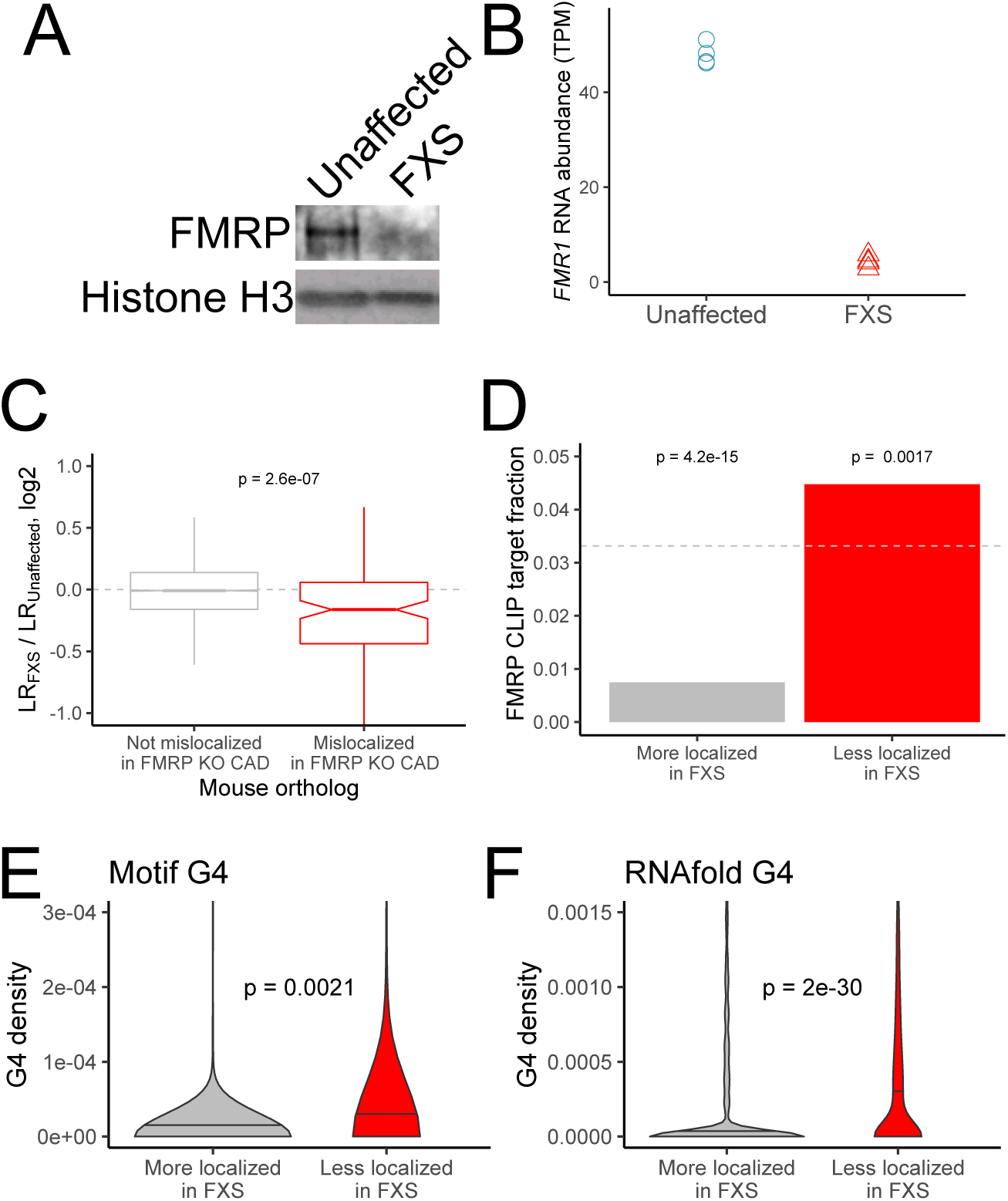
RNA mislocalization in FXS neurons. (A) FMRP protein expression in motor neurons differentiated from iPS cells derived from unaffected and FXS patients. (B) *FMR1* transcript abundance in motor neurons differentiated from iPS cells derived from unaffected and FXS patients. (C) Relative LR values comparing unaffected and FXS motor neurons. Human genes were binned based on whether or not their mouse orthologs were defined as FMRP localization targets (red) or nontargets (gray) in mouse CAD cells. (D) Fraction of human FMRP localization targets (red) and nontargets (gray), as defined in iPS-derived motor neurons, that were previously identified as bound by FMRP in cells (Ascano et al, 2012; Darnell et al 2011). (E-F) G-quadruplex densities in the 3′ UTRs of human FMRP localization targets (red) and nontargets (gray) as defined using a regular expression (E) and RNAfold (F). Y-axis values depict densities as in figures 2C and 2D.

We then defined genes whose localization differed between the FXS and unaffected samples. After quantifying the localization of genes in each sample, we found that very few genes met a reasonable FDR cutoff for mislocalization. This is perhaps due to the inherent variability in the differentiation of iPS cells into motor neurons from prep to prep or the difference in genetic background of the iPS lines adding noise to gene expression measurements (**Fig S3D**). To define mislocalized genes, we therefore used a log2 fold change cutoff to define genes whose neurite-localization decreased in FXS (FMRP targets) or increased in FXS (FMRP nontargets) (**Fig S3F**). We found that FMRP targets were significantly more likely to be directly bound by FMRP in cells as determined by CLIP than FMRP nontargets (**Fig 3D**) (Ascano et al., 2012; Darnell et al., 2011). We also found that FMRP targets had significantly higher densities of G-quadruplexes in their 3′ UTRs than FMRP nontargets, suggesting that this mode of localization target recognition may be conserved between mice and humans (**Fig 3E, F**).

### The RGG domain of FMRP specifically recognizes G-quadruplex sequences

Given our findings that FMRP localization targets were enriched for G-quadruplex sequences, we next sought to determine if FMRP directly interacted with G-quadruplex RNAs over other RNAs in vitro. To this end, we performed RNA Bind-n-Seq (RBNS), a quantitative and unbiased method that defines the specificity of a given RBP or RNA binding domain by incubating purified protein with a diverse pool of billions of random RNA sequences (Dominguez et al., 2018; Lambert et al., 2014; Taliaferro et al., 2016b). In this technique, the input RNA pool and protein-associated RNA are sequenced using high-throughput sequencing. RNA sequences that mediate protein binding are identified as those enriched in the protein-bound RNA relative to the input pool (**Fig 4A**). Importantly, this assay uses a single round of RNA selection and uncovers a continuous range of RBP specificity.

**Figure 4.**
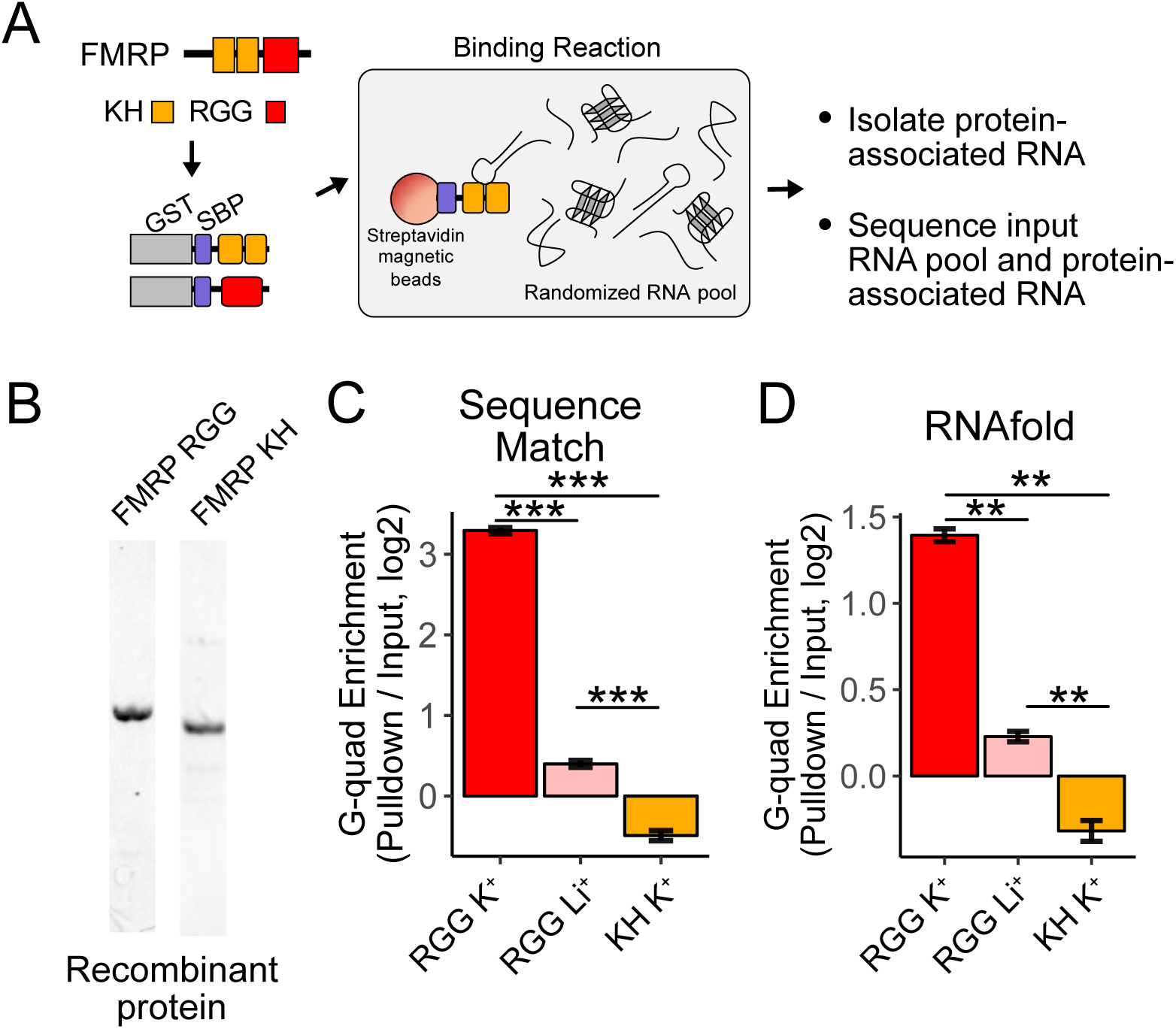
RNA Bind-n-Seq (RBNS) of FMRP domains. (A) Schematic representation of RBNS assay and domain architecture of FMRP. (B) Coomassie stain of purified recombinant FMRP fragments used for RBNS experiments. (C) Enrichments for G-quadruplex RNA sequence motifs in the protein-associated RNA relative to input RNA in the presence of potassium (red) and lithium (pink). Bars represent means of bootstrapped subsets. Error bars represent the standard deviation across subsets. (D) As in C, enrichments for G-quadruplex RNA sequences as predicted by computational RNA folding in protein-associated RNA relative to input RNA. Wilcoxon rank-sum p values < 0.01 are represented as (**), and p values less than 0.001 are represented as (***).

We set out to determine which domains within FMRP, if any, contained the ability to specifically interact with G-quadruplex sequences. FMRP contains two domain types that are known to bind RNA, a pair of KH domains and a single RGG domain (**Fig 4A**). We purified recombinant fragments of mouse FMRP that included either the KH domains or the RGG domain (**Fig 4B**). We then performed RBNS with a pool of random 40mers in the presence of either potassium, which promotes G-quadruplex folding, or lithium, which inhibits G-quadruplex folding (Hardin et al., 1992). For each condition, we sequenced approximately 10 million RNA sequences from both the input and protein-associated RNA pools. G-quadruplex RNA sequences, defined either as matches to specific motifs or through RNA folding predictions, were approximately 3 to 10-fold enriched in the RGG-associated RNA compared to the input pool (**Fig 4C, D**). Furthermore, the addition of lithium drastically reduced this enrichment of G-quadruplex sequences, indicating that the G-quadruplexes must be folded in order to interact with the RGG domain in vitro, as has been previously observed (Menon et al., 2008; Ramos et al., 2003; Schaeffer et al., 2001; Zhang et al., 2014).

The KH domains of FMRP did not specifically interact with G-quadruplex sequences (**Fig 4C, D**), and we were unable to uncover any sequence specificity in the KH-bound RNA. These data suggest that FMRP binds its RNA localization targets through interactions between its RGG domain and target transcripts.

### The RGG domain of FMRP as well as G-quadruplex sequence in target mRNAs are both required for efficient localization of RNAs

To investigate a functional relationship between the RGG domain of FMRP and G-quadruplex sequences within 3′ UTRs of localized mRNAs, we used single molecule FISH (smFISH) to monitor the localization of a reporter transcript in CAD cells that contained either no FMRP, full length FMRP, or a truncation of FMRP lacking the RGG domain. These cells were made by rescuing an FMRP knockout line with GFP, FMRP, or FMRPΔRGG, respectively (**Fig 5A**). The reporter transcript contained the coding sequence of Firefly luciferase followed by the 3′ UTR of the Nol3 gene (Nol3 3′ UTR). This UTR was chosen because we observed in our sequencing data that Nol3 was mislocalized in FMRP knockout CAD cells (LR log2 fold change = −0.9, p = 3e-14). In separate experiments, we also monitored the localization of a version of the reporter transcript in which the only RNAfold-predicted G-quadruplex sequence had been removed (Nol3-G4 3′ UTR) (Lorenz et al., 2011). The smFISH probes used to visualize the two reporters were identical and targeted the coding sequence of the transcript (**Fig 5A**).

**Figure 5.**
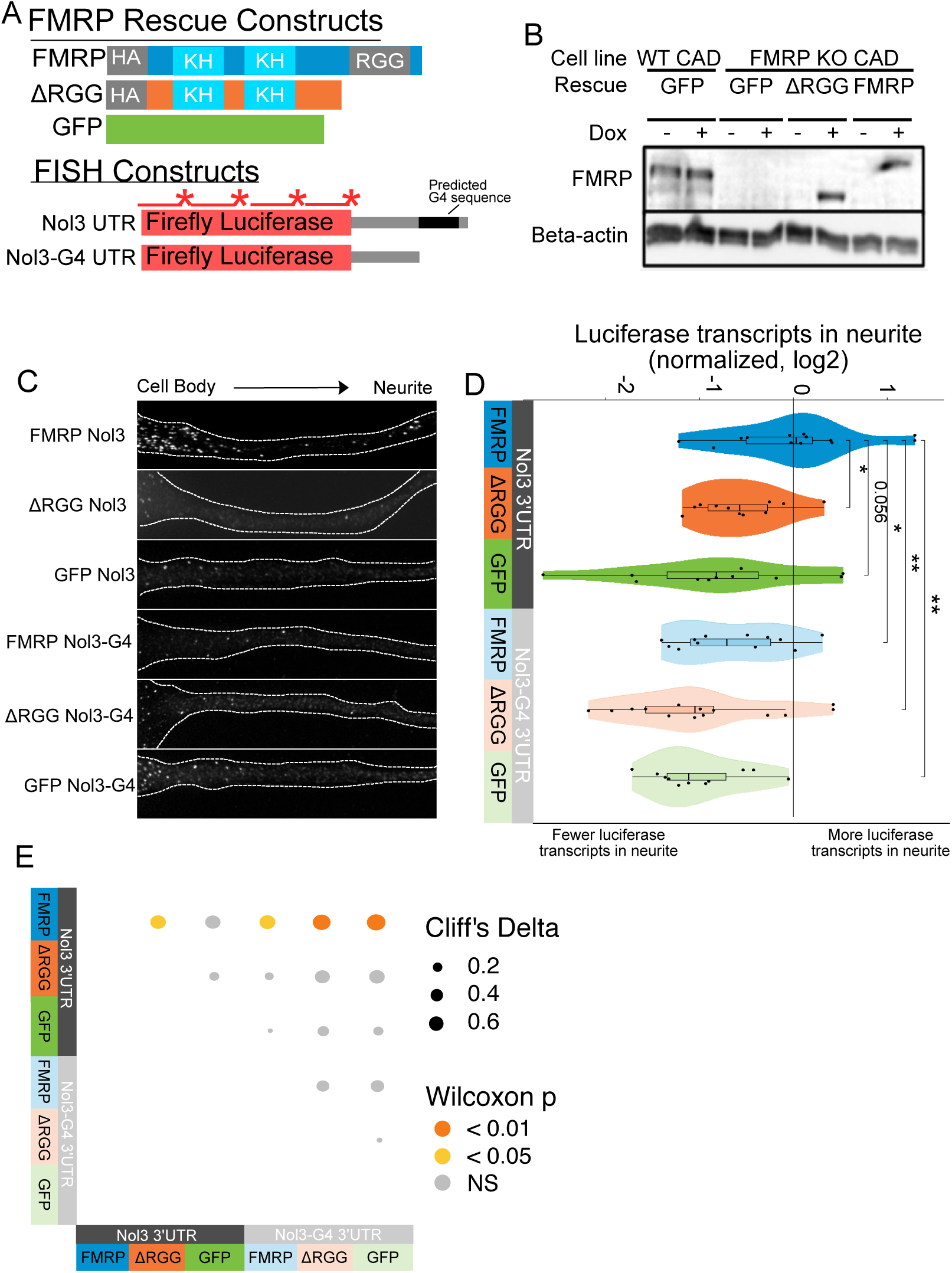
Efficient localization of a reporter transcript requires both a G-quadruplex sequence and the RGG domain of FMRP. (A) FMRP rescue constructs and smFISH reporter constructs used. smFISH probes, represented by red bars with asterisks, hybridize to the ORF of both reporter constructs. (B) Expression of FMRP rescue constructs in FMRP knockout-rescue CAD cells. (C) Representative Firefly Luciferase transcript FISH images. Images are oriented such that the cell body is toward the left. (D) Quantification of FISH images. For all conditions, the number of reporter transcripts in the neurite was calculated using FISH-quant. Each dot in each condition represents a single cell. Transcript counts were normalized to those observed in the full length FMRP, Nol3 UTR condition. P values were calculated using a Wilcoxon rank-sum test. (E) Summary statistics of results in D. Effect sizes are represented using Cliff’s delta.

Both the FMR1 rescue constructs and the reporter transcripts were integrated into the genome of the CAD cells using Cre-lox recombination into a single genomic site (Khandelia et al., 2011). Using this strategy, cell pools were virtually homogeneous for integration of the rescue constructs, and the expression of both the rescue constructs and the reporter transcripts were controllable with doxycyline. Protein levels were assayed via western blot with or without doxycycline induction to confirm expression of rescue constructs. We found that the expression level of FMRP in the rescued cells was similar to the expression of FMRP in wildtype cells (**Fig 5B**). We then monitored the efficiency of reporter transcript localization to neurites by smFISH and quantified the results using FISH-quant (Mueller et al., 2013). We observed that the reporter transcript that contained a G-quadruplex sequence in its 3′ UTR was efficiently localized to neurites, but only in cells expressing full length FMRP (FMRP, Nol3 3′ UTR), and not in cells either completely lacking FMRP (GFP, Nol3 3′ UTR) or in cells expressing only a truncated form of FMRP (ΔRGG, Nol3 3′ UTR) (**Fig 5C-E**). Removal of the G-quadruplex sequence (Nol3-G4 3′ UTR) resulted in less efficient transcript localization regardless of the FMRP status of the cell (**Fig 5C-E**). These results are consistent with both the RGG domain of FMRP and a G-quadruplex sequence within the target transcript being required for efficient localization to neurites.

To probe the functional relationship between the RGG domain of FMRP and G-quadruplex sequences transcriptome-wide, we fractionated the FMRP rescue lines into soma and neurite fractions and analyzed RNA from those fractions using high-throughput sequencing. We observed enrichments for mRNAs encoding ribosomal proteins and electron transport chain components in the neurites of all samples, indicating that fractionation of the cells was successful (**Fig S4A**). Transcriptome-wide measures of RNA localization, as measured by LR values of all genes in the GFP and ΔRGG lines were strongly correlated. This is consistent with neither of these lines containing a localization-competent form of FMRP. Full length FMRP rescue samples were outliers in this analysis compared to the GFP and ΔRGG samples, perhaps due to the active localization regulatory activity of full length FMRP (**Fig 6A, Tables S3-5**).

**Figure 6.**
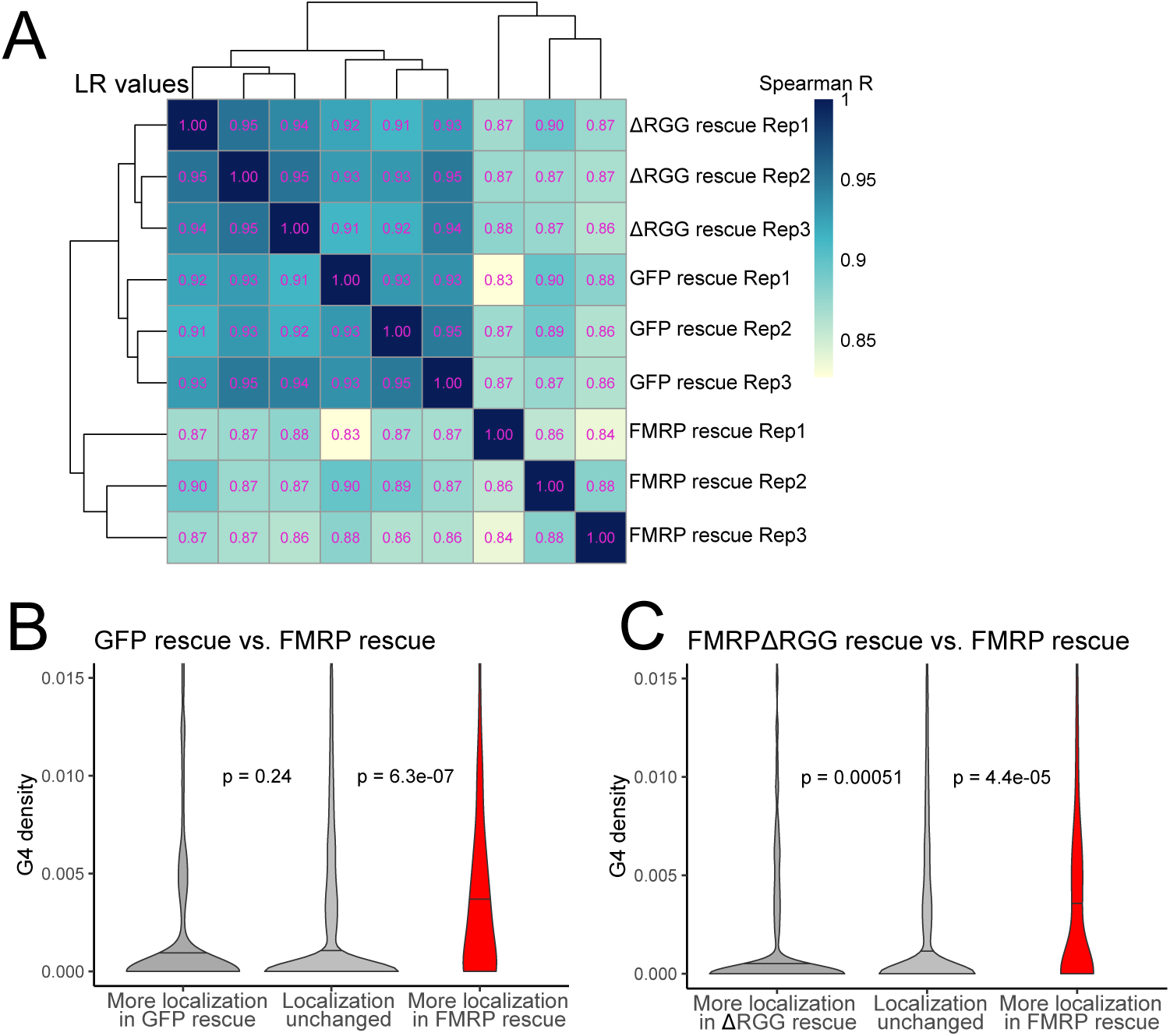
The RGG domain of FMRP is required for the localization of G-quadruplex-containing transcripts transcriptome-wide. (A) Correlation of LR values for all expressed genes in FMRP rescue samples. (B) G-quadruplex density, as measured by RNAfold, in the 3′ UTRs of transcripts that were more localized in the GFP rescue cells (dark gray, left) or full-length FMRP rescue cells (red, right). P values were calculated using Wilcoxon rank-sum tests. As in figure 2D, values represent the number of guanosine residues participating in quadruplex sequences per nt. (C) As in (B), but comparing transcripts that were more localized in the FMRP RGG truncation rescue (dark gray, left) or the full length FMRP rescue (red, right).

We then compared changes in localization across samples by looking at changes in LR values for all genes. We found that comparing LR values in the full length FMRP rescue to either the GFP rescue or ΔRGG rescue produced highly concordant results (**Fig S4B**), consistent with both comparisons involving one functionally active FMRP condition and one functionally inactive FMRP condition. These comparisons were also strongly positively correlated with our original CAD knockout experiment in which FMRP null and wildtype cells were compared (**Fig S4B**).

We then compared the 3′ UTR G-quadruplex densities of genes that were differentially localized between given rescue conditions. When we compared the GFP rescue samples and full length FMRP rescue samples, we found that genes that were significantly more localized in the FMRP rescue cells contained significantly more G-quadruplex sequences in their 3′ UTRs than expected, consistent with our previous results (**Fig 6B, 2D**). When we performed the same analysis comparing the ΔRGG rescue samples to the full length FMRP rescue samples, we again found that genes whose localization was significantly more efficient in the full length FMRP samples contained significantly more 3′ UTR G-quadruplex sequences than expected (**Fig 6C**). This suggests that, transcriptome-wide, the RGG domain of FMRP is important for the neurite-localization of RNA molecules that contain G-quadruplex sequences. Consistent with this, we observed no relationship between the G-quadruplex content of an RNA and its differential localization in the GFP rescue and ΔRGG rescue samples (**Fig S4C**).

### The translational regulatory targets and RNA localization targets of FMRP are distinct pools of transcripts

FMRP is known to regulate mRNA metabolism at the level of RNA localization and translation (Darnell et al., 2011; Dictenberg et al., 2008). We wondered whether the transcripts targeted by these activities were largely the same or distinct. To address this question, we identified FMRP translational regulatory targets by performing ribosomal profiling on wildtype and FMRP null CAD cells (Ingolia et al., 2009). We observed that our ribosome protected fragments displayed strong enrichments for coding regions as well as a strong 3 nucleotide periodicity, indicating that the ribosome profiling data was of high quality (**Fig S5A-D**).

We used the software package Xtail to identify transcripts that displayed significantly different ribosome occupancies in wildtype and FMRP null CAD cells (**Fig S5E, Table S6**) (Xiao et al., 2016). Previous studies had reported that FMRP predominantly negatively regulates translation by inducing ribosome stalls (Darnell et al., 2011). Since stalls may result in ribosome pileups and increased ribosome densities, we reasoned that FMRP translation targets should have a lower ribosome occupancy in FMRP null cells, even though their rate of translation and protein output is increased in the absence of FMRP (**Fig 7A**). Consistent with this, we found genes that displayed reduced ribosome occupancy in FMRP null cells compared to wildtype cells were significantly more likely to have been identified as FMRP CLIP targets than expected (**Fig 7B**) (Ascano et al., 2012; Darnell et al., 2011).

**Figure 7.**
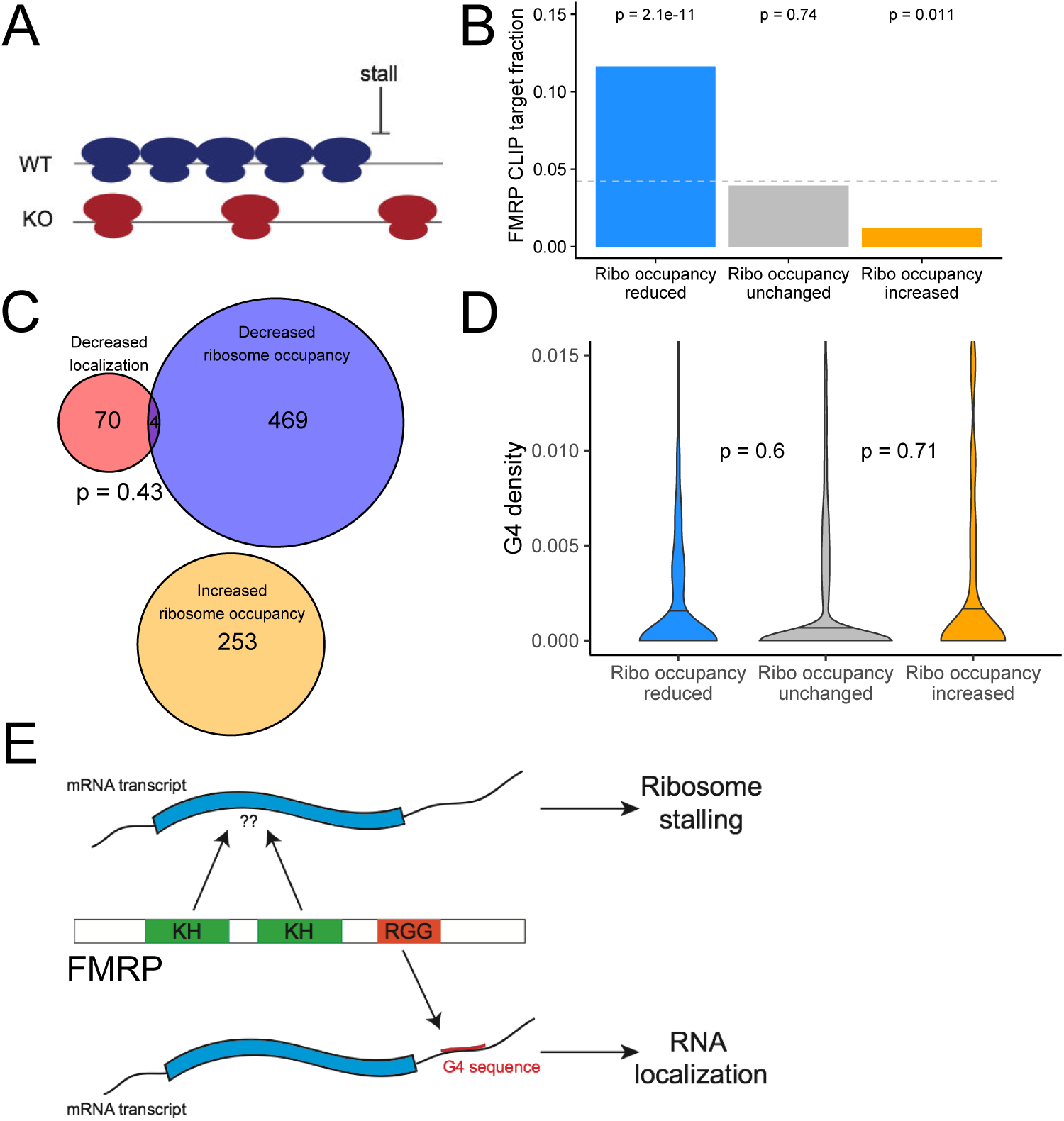
The translational regulation targets of FMRP are distinct from its RNA localization targets and have no relationship to G-quadruplex sequence density. (A) Schematic of how the removal of FMRP, which is known to cause ribosome stalls, may affect overall ribosome density on a transcript. (B) Fraction of genes whose ribosome occupancy significantly decreased (blue) or increased (orange) in the FMRP null samples that were previously identified as bound by FMRP in cells. The null expectation is represented by a gray dotted line. P values were calculated using a hypergeometric test. (C) Overlap of FMRP localization targets (red) and translational repression targets (blue). P values were calculated using a Binomial test. (D) G-quadruplex density, as determined by RNAfold, for the 3′ UTRs of transcripts whose ribosome occupancy was decreased (blue) or increased (orange) in the FMRP-null samples. P values were calculated using a Wilcoxon rank-sum test. (E) Model for how FMRP recognizes its targets. RNA localization targets are recognized through the interaction of the RGG domain and a G-quadruplex sequence in the 3′ UTR of the transcript. Translational regulation targets are recognized through the interaction of one or both KH domains with an unknown transcript feature, or perhaps through interaction with ribosomes.

We found a significant but weak correlation between the mislocalization (as measured by changes in LR) and translational repression (as measured by changes in ribosome occupancy) of transcripts due to FMRP (R = 0.08, p = 4.1e-16). However, when we looked at the transcripts that were statistically significantly affected in the two conditions, we found no significant overlap between the translational regulatory targets and the RNA localization targets of FMRP, suggesting that the protein exerts these activities on largely distinct pools of transcripts (Fig 7C). While the localization targets of FMRP were enriched for genes that regulate transport in cells (**Fig S1G**), the translational regulatory targets were enriched for genes that regulate RNA metabolism and for those that regulate synaptic activity (**Fig S5F**).

We also found no relationship between the 3′ UTR G-quadruplex content of a transcript and whether or not it was translationally regulated by FMRP (**Fig 7D**). Identifying transcript features associated with translational regulation by FMRP proved difficult as the transcript features most strongly associated with FMRP translation targets were the GC content and length of the transcript’s CDS (**Fig S5G, H**). Translational regulatory targets of FMRP were strongly enriched for having GC-poor, long CDS sequences. These effects are difficult to disentangle, however, as transcript GC content and length are correlated with each other in multiple species (Marín et al., 2003) as well as with a variety of other metrics, including conservation, expression, and RNA secondary structure.

Given that the transcript targets and modes of target recognition for the two activities of FMRP appear to be distinct, we propose a model in which the recognition of a G-quadruplex sequence within the 3′ UTR of a transcript by the RGG domain of FMRP is associated with localization of the transcript. Conversely, the recognition of some as-yet unidentified transcript feature by one or both of the KH domains of FMRP is associated with translational regulation (**Fig 7E**). This transcript feature, if it exists, has been difficult to identify as both bioinformatic sequence analysis of the translational regulation targets of FMRP and the RNA sequences specifically bound by the KH domains of FMRP in the RBNS experiment yielded no strongly enriched motifs.

This model is supported by three previously reported observations. First, the RGG domain of FMRP is dispensable for the interaction of the protein with polyribosomes (Darnell et al., 2005b). Second, the FXS-associated FMR1 mutation I304N, which is in one of the KH domains, abolishes polyribosome association (Feng et al., 1997). Third, G-quadruplex sequences are unable to compete with polysomes for binding to FMRP (Darnell et al., 2005a). Taken together with the data presented here, this supports the idea that the dual functionalities of FMRP are driven by distinct binding capabilities of its different RNA binding domains.

## DISCUSSION

Hundreds to thousands of transcripts are asymmetrically distributed within cells, yet the mechanisms that regulate trafficking are known for only a handful (Cajigas et al., 2012; Gumy et al., 2011; Lécuyer et al., 2007; Taylor et al., 2009; Wang et al., 2012). In this study we have used genome engineering, subcellular fractionation, and high-throughput sequencing to identify transcripts that depend on FMRP for transport in neuronal cells. These FMRP localization targets displayed a large enrichment for G-quadruplex sequences, particular in their 3′ UTRs, suggesting that FMRP specifically recognizes these sequences to target them for transport. G-quadruplex sequences were not enriched in the translational regulatory targets of FMRP as identified by ribosome profiling. In fact, no recognizable sequence motif was strongly enriched in the translational regulatory targets beyond a negative correlation with GC content. These apparent dual modes of target recognition and the lack of clear specificity in the translational regulatory targets may explain some of the previously reported conflicting results in regards to the prefered RNA binding sites of FMRP (Anderson et al., 2016).

FMRP has long been known to bind to G-quadruplex sequences (Anderson et al., 2016; Darnell et al., 2001; Suhl et al., 2014), and G-quadruplex sequences have been shown to be sufficient for transport to neurites (Subramanian et al., 2011). It is important to note that we cannot formally prove that the G-quadruplex sequences enriched in FMRP localization targets are actually folded in cells. Recent reports have demonstrated that the majority of RNA sequences that have the ability to fold into G-quadruplex structures in vitro may be generally unfolded in cells (Guo and Bartel, 2016). Still, previous reports and our data demonstrate that the G-quadruplexes must be folded in order to be efficiently bound by FMRP (Subramanian et al., 2011), and G-quadruplex sequences have been reported to be under purifying selection (Lee et al., 2019).

No clear links have yet been found between the mislocalization of a specific transcript or transcripts in FXS neurons and FXS phenotypes. The relative contribution of the RNA localization and translational regulation activities of FMRP to FXS phenotypes similarly remain unclear. Further work is needed to more fully understand mechanistic links between RNA localization and neurological diseases including FXS.

For the vast majority of localized transcripts, in neurons and elsewhere, the RBP/RNA interactions that mediate transport are completely unknown. Classically, experiments designed to address these questions have focused on one transcript at a time and take the approach of systematic deletions and/ or mutations to identify minimal sequence elements that regulate localization (Ferrandon et al., 1994; Jambor et al., 2014; Kim-Ha et al., 1993; Kislauskis et al., 1994). While these experiments have been successful and provided key insights for the field, they are quite labor-intensive, and it is difficult to derive general rules that govern RNA localization from isolated, single-transcript examples. RBP-perturbation experiments followed by transcriptomic profiling have been highly successful in elucidating mechanisms that underlie other RNA metabolic processes, including pre-mRNA processing, translation, and decay (Iwasaki et al., 2016; Mukherjee et al., 2011; Wang et al., 2012). By extending these techniques to the study of RNA localization, we envision this study as providing a framework for the investigation of other RBPs and their effects on RNA trafficking. Given that RNA mislocalization has been associated with a range of neurological diseases (Wang et al., 2016), these studies can lay groundwork that links molecular defects to observed phenotypes.

## METHODS

### Construction of an FMRP-null CAD line

The guide RNA sequence AAATTATCAGCTGGTAATTT was cloned into pX330 (gift of F. Zhang) using the BbsI sites. This plasmid was then transfected into CAD cells using Lipofectamine LTX. Forty-eight hours after transfection, the cells were sorted to single cells using a flow cytometer and allowed to grow without interference for two weeks. Clones were screened for FMRP expression using a polyclonal FMRP antibody (MBL International RN016P). Lesions at the locus of cutting were identified by PCR using genomic DNA to amplify the locus followed by Topo cloning of the PCR fragment and Sanger sequencing of individual bacterial colonies.

### Sequencing of subcellular transcriptomes from wildtype and FMRP null CAD cells

CAD cells were grown in DMEM/F12 (Gibco 11320033) supplemented with 10% FBS at 37C in 5% CO2. To fractionate CAD cells into soma and neurite fractions, CAD cells were plated on porous, transwell membranes that had a pore size of 1.0 micron (Corning 353102). One million CAD cells were plated on each membrane. These membranes fit in one well of a six well plate, and six membranes were combined to comprise one single preparation. After allowing the cells to attach, the cells were induced to differentiate into a more neuronal state through the withdrawal of serum. Cells were then differentiated on the membranes for 48 hours.

To fractionate the cells, the membranes were washed one time with PBS. One mL of PBS was added to the top (soma) side of the membrane. The top side of the membrane was then scraped repeatedly and thoroughly with a cell scraper and the soma were removed into an ice-cold 15 mL falcon tube. After scraping, the membranes were removed from their plastic housing with a razor blade and soaked in 500 µL RNA lysis buffer (Zymo R1050) at room temperature for 15 min to make a neurite lysate. During this time, the 6 mL of soma suspension in PBS was spun down and resuspended in 600 uL PBS. 100 µL of this soma sample was then carried forward for RNA isolation using the Zymo QuickRNA MicroPrep kit (Zymo R1050). In parallel, RNA was also isolated from the 500 uL neurite lysate using the same kit. Typically, the neurite RNA yield from the six combined membranes was 500-1000 ng. The efficiency of the fractionation was assayed by Western blotting using anti-beta-actin (Sigma A5441, 1:5000) and anti-Histone H3 (Abcam ab10799, 1:10000) antibodies.

PolyA-selected, stranded libraries were constructed using 25 ng total RNA and an Illumina Neoprep library preparation system. The resulting libraries were then sequenced on an Illumina NextSeq 500 sequencer using paired-end, 75 bp reads. Approximately 40 million read pairs were obtained per sample. Each condition (wildtype and FMRP null; soma and neurite) was prepared and sequenced in triplicate for a total of 12 samples.

### Quantification of RNA localization from CAD subcellular sequencing data

Transcript abundances were calculated from read files using kallisto (CAD Fmr1 knockout) (Bray et al., 2016) or using salmon v0.11.1 (CAD Fmr1 rescue samples, FXS iPS samples) (Patro et al., 2017) with the options --seqBias and --gcBias. Transcript abundances were then collapsed to gene abundances using txImport (Soneson et al., 2015). LR values for genes were calculated as log2 of the ratio of neurite/soma normalized counts for a gene produced by DESeq2 using the salmon/tximport calculated counts (Love et al., 2014). Genes with less than 10 counts in any sample were excluded from further analyses. To identify genes whose LR values significantly changed across conditions, the Xtail software package was used (Xiao et al., 2016). Xtail is designed to process ribosome profiling data and identify genes whose ribosome occupancy changes across conditions. However, since ribosome occupancy is simply a ratio of expression count values (ribosome footprints / RNAseq), we reasoned that since LR is also a ratio of expression count values (neurite / soma), Xtail could be used to identify genes whose LR value changed across conditions. For further analyses of this data, mislocalized genes were defined as those with a FDR of less than 0.01.

### Identification of transcript features associated with mislocalized genes

To ask if previously identified FMRP-interacting transcripts were enriched in our set of mislocalized genes, we defined FMRP-interacting transcripts as the intersection of RNA targets identified in two earlier studies (Ascano et al., 2012; Darnell et al., 2011) as was previously done (Suhl et al., 2014).

To identify 6mer sequences enriched in the 3′ UTRs of mislocalized genes, we calculated the abundance of all 6mers in the UTRs of mislocalized genes and, as a control, in the UTRs of expressed but non-mislocalized genes. These abundances were then compared using a Fishers exact test, and the resulting p values were corrected for multiple hypothesis testing using the Benjamini-Hochsberg method.

To computationally identify predicted G-quadruplex sequences, we used two approaches. First, to identify WGGA sequences separated by variable length linkers, we searched sequences using the regular expression (?=([AU]GGA(.{0,6})[AU]GGA(. {0,6})[AU]GGA(.{0,6})[AU]GGA)). Alternatively, we analyzed sequences using a sliding window approach and RNAfold (Lorenz et al., 2011). The window was 80 nt wide and slid 20 nt at a time. The sequence within each window was analyzed using RNAfold and the −g option. Nucleotides with a ‘+’ for their predicted structure status were designated as nucleotides participating in a quadruplex.

For analyses involving experimentally defined G-quadruplex sequences, data was taken from previously published experiments using RNA purified from mouse embryonic stem cells that utilized RT stops that occurred with differential frequency in G-quadruplex promoting (potassium) conditions and G-quadruplex inhibiting (lithium) conditions (Guo and Bartel, 2016). The data taken from this paper contained RT stop sites, the 60 nt upstream of the stop site, and the frequency at which those sites were observed in the potassium and lithium conditions. RT stops representing G-quadruplexes were defined as those which were observed at least 5 times more frequently in the potassium data than in the lithium data. For those sites, the G-quadruplex region was defined as the 60 nt immediately upstream of the stop.

Cliff’s delta values representing the effect sizes associated with each transcript feature and its association with FMRP-mediated localization were calculating using the orddom package in R.

### G-quadruplex conservation analyses

To ask if G-quadruplexes were positionally conserved, PhyloP scores were calculated for all G nucleotides in all 3′ UTRs. If the G nucleotides were predicted by RNAfold to be participating in quadruplexes as discussed above, their scores were put in one bin. If they were not predicted to be participating in quadruplexes, they were put in another bin. The scores of the G nucleotides in the two bins were then compared.

### FXS iPSC line generation and iPSC culture

iPSCs were generated by transducing control and FXS patient fibroblasts with Sendai virus from the Cytotune 2.0 Reprogramming Kit (Life Technologies) per the manufacturer’s protocol. Briefly, early passage fibroblasts were cultured in fibroblast medium (10% ES-qualified FBS, 0.1 mM NEAA, 55µM β-mercaptoethanol, high glucose DMEM with Glutamax) for two days. On day 0, fibroblasts were transduced with Sendai virus encoding KOS, hc-Myc, hKlf4, each at an MOI of 5. Cells were fed with fibroblast medium every other day for one week. On day 7, cells were passaged onto vitronectin (Life Technologies) coated dishes at a density of 250,000 to 500,000 cells/well. Beginning on day 8, cells were fed every day in Essential 8 medium (Life Technologies) and colonies began to emerge within another 7-10 days. Individual iPSC colonies were manually picked and transferred to a dish coated with either vitronectin or Matrigel (BD) and expanded as clonal lines. All iPSC lines were karyotyped (WiCell) and characterized for expression of markers of pluripotency using immunofluorescence and RT-PCR. Additionally, we confirmed loss-of-function of FMR1 in our FXS patient iPSC lines by taqman assays for FMR1 mRNA and western blotting for FMRP. iPSCs were maintained on Matrigel coated dishes and fed every day with complete mTesR1 medium (Stem Cell Technologies). iPSCs were passaged every 5-7 days using ReLeSR (Stem Cell Technologies).

### FXS iPS line differentiation into motor neurons

Motor neurons were generated using an improved, suspension culture based protocol based on previously published Dual-Smad inhibition approaches (Chambers et al., 2009; Du et al., 2015; Reinhardt et al., 2013). iPSCs were grown as clusters in suspension, and differentiated in a set-wise fashion as outlined in Supp. Fig.3a. Briefly, clusters were exposed to Smad and ROCK inhibition and WNT activation during the first 6 days, followed by activation of the retinoic and sonic hedgehog pathways. At day 10 the clusters were dissociated, and single cells were plated on membranes coated with Matrigel (Corning) in maturation media until d18-21 when MNs were analyzed.

### Fractionation of iPS neurons into soma and neurite fractions

Generally, iPS neurons were fractionated in a very similar method as the CAD cells. However, since the iPS neurons usually contained less material, instead of combining 6 wells together into one prep as was done for the CAD cells, between 6 and 12 wells were combined together. After mechanical fractionation by scraping into soma and neurite fractions and RNA isolation using the Zymo Quick RNA Microprep Kit, between 100 and 500 ng of neurite RNA was obtained.

Fractionation efficiency was monitored by dot blot using the beta-actin and histone H3 antibodies described above. PolyA-selected, stranded RNAseq libraries were constructed using the Kapa mRNA Hyperprep kit, 100 ng of total RNA, and 15 PCR cycles of library amplification. Each condition (FXS and unaffected; soma and neurite) was fractionated, prepared, and sequenced with 4 replicates for a total of 16 samples. Samples were sequenced using paired-end, 150 nt sequencing on an Illumina NovaSeq. Approximately 30 million read pairs were obtained per sample.

### Analysis of iPS neuron subcellular transcriptome data

Raw sequencing data from the iPS soma and neurite samples were analyzed using salmon and tximport as described above. After analysis of changes in RNA localization between FXS and unaffected samples using Xtail, very few genes met a reasonable FDR threshold for being significantly different between conditions. This may be a reflection of the increased variance in expression values in these samples relative to the CAD samples, perhaps due to slight variations between preps in the differentiation from iPS cells into motor neurons. For the analysis of G-quadruplex densities in mislocalized genes, mislocalized genes were identified as those who had a log2 fold change in LR across conditions (FXS vs unaffected) of at least 0.25.

### RNA Bind-n-Seq protein purification and sequence analysis

Rosetta E. coli cells were transformed with pGEX6 bacterial expression constructs (GE Healthcare) harboring GST-SBP (streptavidin binding peptide)-RGG or the two KH domains from mouse FMRP. The RGG domain was defined as residues 478-614 and the KH domains were defined as residues 222-315 from mouse FMRP. Cultures were grown in LB broth until the optical density reached ∼0.8. Cultures were brought to 16°C and induced with IPTG overnight. Cells were then harvested and lysed in lysis buffer (25mM Tris-HCl, 150mM NaCl, 3mM MgCl2, 1% Triton X-100, 500units/per 1L culture Benzonase Nuclease, 20mg/L Lysozyme, Pierce Protease Inhibitor tablets). Cell pellets were then sonicated, incubated for 30min on ice, and centrifuged at 12500 rpm for 40 minutes. Supernatants were passed through a 0.45 micron filter and GST-tagged proteins were purifed using GST-trap FF columns (GE). Samples were further subjected to size exclusion chromatography (Superdex 200 Increase 10/300 GL column (GE)). Pure fractions were pooled together and concentrated by centrifugation (Amicon Ultra-0.5mL Centrifugal Filter Units). Purity was assessed by PAGE and Coomassie stain.

The RNA Bind-n-Seq assay was performed similar to previously described (Dominguez et al., 2018). Briefly, RNA with a random region of 40nt was prepared by in vitro transcription. The RNA pool was resolved by PAGE and purified. Concentrated RNA was heated at 85C for 5 min in the presence of 150mM KCl or LiCl and allowed to cool for at least 3 min to allow for folding. Binding reactions were performed as previously described (Dominguez et al., 2018). All wash and binding buffers contained either 150mM KCl or LiCl. Protein-RNA complexes were eluted, RNA isolated and prepared for sequencing as previously described (Dominguez et al., 2018).

Sequence enrichments in RNA Bind-n-Seq data were obtained in three ways. First, kmer frequencies (k=3-6) were computed in protein-bound libraries and compared to kmers in the input library. For each kmer, the ratio of its frequency in the protein-bound library to the input library results in an enrichment or “R” value for every kmer, as was previously reported (Dominguez et al., 2018). To identify G quadruplex sequences, we determined the fraction of protein-bound reads with matches to a variety of G quadruplex motifs (e.g. GGGN1-7GGGN1-7 GGGN1-7GGG) and compared that to the number of input reads with the same motif. Lastly, we used RNAfold version 2.4.11 (Lorenz et al., 2011) to predict G quadruplexes (RNAfold −g option) in the 40 nucleotide randomized regions of input and bound RNAs. As with the pattern match approach, the ratio of protein-bound to input reads predicted to fold into G quadruplexes was computed. In each case reads were subsampled for estimates. Error bars shown in figures represent standard deviation from mean across subsamples.

### Construction of an FMRP-null CAD line with single LoxP cassette

CAD LoxP cell lines were received from the lab of Eugene Makeyev (Khandelia et al., 2011). This line contains a single loxP cassette in the genome of the cells to facilitate recombination of exogenous constructs. A knockout insertion cassette was introduced to exon 6 of the endogenous Fmr1 gene via CRISPR-Cas9 technology to generate CAD LoxP FMRP KO cells. This cassette includes stop codons in all frames as well as a strong polyadenylation signal. 300 nt of homology on either side of the gRNA cut site was included to facilitate homology directed repair. A G418 resistance gene was included to select for integration. pX330 was used to express Cas9 and the guide RNA sequence AAATTATCAGCTGGTAATTT. Following selection with G418, cells were sorted to single cells using a flow cytometer and allowed to expand for two weeks. Clones were screened for insertion of the cassette by PCR as well as FMRP mRNA and protein depletion determined by qPCR, RNA-seq and Western Blot Analysis with a monoclonal FMRP antibody (Proteintech 66548). Lesions at the locus of cutting were identified by PCR using genomic DNA to amplify the locus followed by Topo cloning of the PCR fragment and Sanger sequencing of individual bacterial colonies.

### Expression of Fmr1 rescue constructs in CAD cells

CAD LoxP FMRP KO cell lines were plated in 12-well plates at 1.0 to 1.5 × 105 cells per well in an DMEM:F12 supplemented with 10% Equafetal 12 to 18 hours before transfection. Cells were then co-transfected with a derivative of pRD-RIPE (containing one of three Fmr1 Rescue cDNAs and Firefly Luciferase fused with one of two 3’UTRs) blended with 1% (wt/wt) of a Cre-encoding plasmid (pCAGGS-Cre). To transfect one well of a 12-well plate, 0.5 μg of purified plasmid was mixed with 2 μL Lipofectamine LTX reagent, 1 uL PLUS reagent and 50 μL Opti-MEM following the manufacturer’s protocol. Cells were incubated with the transfection mixtures overnight, the medium was changed and the incubation continued for another 24 hours before addition of puromycin (5ug/ml). Incubation with puromycin continued for several days until untransfected control wells were depleted of cells. The cultures with stably integrated constructs were expanded in a full growth medium. Lysates were collected following 2ug/mL Doxycycline induction to assay construct expression via western blot.

### Immunoblot of FMRP rescue constructs

Expression of rescue constructs in the CAD FMRP KO cell lines were determined by Western blot analysis. Cells with integrated rescue constructs were plated in 6-well plates at 2-3 × 105 cells per well in duplicate. Each cell line was grown in full growth media or full growth media supplemented with 2ug/mL doxycycline for 48 hours. Lysates were collected with 200uL RIPA buffer, incubated on ice for 15 minutes then passed through a 20G needle 20 times. Lysates were stored at −80°C. Denatured lysates were separated by PAGE on 4%–12% Bis-Tris gradient gels (Invitrogen) along with Spectra Multicolor Broad Range (Fisher Scientific) using MOPS SDS NuPAGE Running Buffer (Invitrogen) and NuPAGE LDS Sample Buffer (Invitrogen) on ice at 200 V for 1 hour. Gels were transferred to PVDF membranes (Millipore) using NuPAGE Transfer Buffer (Invitrogen) at 15 V for 55 minutes before being washed three times, 5 minutes each with PBST at room temperature (RT) with agitation and blocked with 5% Milk Powder (Sigma) in PBST for 1 hr at RT with agitation. Blots were again washed 3 times, 5 minutes each in PBST at RT with agitation before being incubated in 1:5000 FMRP mouse monoclonal (Proteintech) primary antibody in 2% Milk Powder in PBST overnight at 4C with agitation or with 1:10,000 Histone H3 mouse monoclonal antibody (Abcam) in 2% milk powder in PBST for 2 hours at room temperature with agitation. Blots were washed 3 times, 5 minutes each at RT with agitation and then transferred to Anti-mouse IgG HRP-conjugated secondary antibody in 2% Milk Powder in PBST and incubated at RT for 2 hours. Blots were washed again as previously described and visualized using WesternBright HRP Substrate Kit (Advansta).

### smFISH of reporter transcripts

CAD LoxP FMRP KO cells with Integrated Rescue constructs were plated on PDL coated glass coverslips (neuVitro) within 12-well plates at approximately 2.5 × 104 cells per well in full growth media. Cells were allowed to attach for 2 hours before changing to serum depleted media supplemented with 2ug/mL doxycycline. Cells were allowed to express constructs and differentiate for 48 hours. Cells were washed once with PBS before being fixed in 3.7% formaldehyde in PBS for 10 minutes at room temperature then washed twice with PBS before permeabilization with 70% Ethanol at 4°C for 6-8 hours. Cells are incubated in smFISH Wash Buffer at room temperature for 2-5 minutes. Per hybridization reaction, 2 uL of Stellaris FISH Probes, Firefly Luc with Quasar® 670 Dye were added to 200uL of Hybridization Buffer. An empty opaque tip box is prepared with wet paper towels below with parafilm covering the top. Hybridization buffer with probes (200uL per reaction) was “beaded” on top of the parafilm and the PDL coated glass coverslips were placed cell side down onto “beaded” buffer. The entire tip box is incubated at 37°C over night. Glass coverslips were transferred to fresh 12-well plates cell side up and incubated with smFISH Wash Buffer at 37°C in the dark for 30 minutes. Buffer was then replaced with fresh smFISH Wash Buffer supplemented with 100ng/mL DAPI and incubated in the dark at 37°C. DAPI stain was replaced with fresh smFISH Wash Buffer and incubated for 5 minutes at room temperature. Coverslips were then mounted onto slides with 6uL Fluoromount G and sealed with nail polish. Slides were imaged on a widefield DeltaVision Microscope with consistent laser intensity and exposure times across samples.

### smFISH quantification

Experimental parameters within FISH-quant were set to match widefield DeltaVision microscope and the Stellaris FISH probes (Tsanov et al., 2016). The excitation and emission wavelengths used were 547nm and 583nm respectively which were recorded by a 60X objective with 1.4 numerical aperture with 1.33 refractive index oil. This was captured by a camera such that pixel sizes in the xy were 98.4nm wide and z pixels a 398nm wide. Cell outlines were drawn using cell autofluorescence in the FITC channel where somas and neurites were treated as separate cells. Images were first filtered with a 3D 2x gaussian method using the default parameters. Pre-detection was conducted, and spots were fit using FISH-quant default settings and a pixel-intensity threshold set to overestimate detected spots. All thresholding parameters were set tightly around each mean population to exclude gross outliers. Averaged detected spots were inspected for spherical character to ensure quality of detection. These detection settings were the same for every soma and neurite imaged.

### Ribosome footprinting of CAD cells

CAD cells were grown in the absence of serum for 48 hours to induce neuronal differentiation. Ribosome footprint libraries and companion RNAseq libraries were then produced using the Truseq Ribo Profiling Kit from Illumina according to the manufacturer’s instructions. Libraries were sequenced on an Illumina NextSeq sequencer, with about 40 million reads being obtained per sample. Three techincal replicates were performed per condition (WT and knockout; ribosome protected fragments (RPF) and RNAseq) for a total of 12 samples.

Gene level abundance estimates from the RPF and RNAseq data were performed using kallisto and tximport as described above (Bray et al., 2016). Ribosome occupancy values and their changes between wildtype and knockout cells were then calculated using Xtail (Xiao et al., 2016).

## DATA AVAILABILITY

Raw sequencing data and processed files are available through the Gene Expression Omnibus, accession GSE137878.

## ACKNOWLEDGEMENTS

We thank Christopher Burge for initial support with this study and members of the Taliaferro lab for helpful discussion.

This work was funded by the RNA Bioscience Initiative at the University of Colorado Anschutz Medical Campus (JMT), a Webb-Waring Early Career Investigator Award from the Boettcher Foundation (AWD-182937) (JMT), NIH1R35GM133885 (JMT), NIHDK120444 (HAR), NIHAI140044 (HAR), NIHDK118803-01A1 (LIH), and NIH1R01MH109026 (GJB). It was further supported by a Pre-doctoral Training Grant in Molecular Biology (NIH-T32-GM008730) (RG), a Guild ACORN Postdoctoral Fellowship (LIH), and grants from the Colorado Clinical and Translational Science Institute (HAR), the Children’s Diabetes Foundation (HAR), a new investigator award from the NIH/HIRN consortium (HAR), the Culshaw Junior Investigator Award (HAR), and the Juvenile Diabetes Research Foundation (HAR).

## SUPPLEMENTARY FIGURES

**Figure S1.**
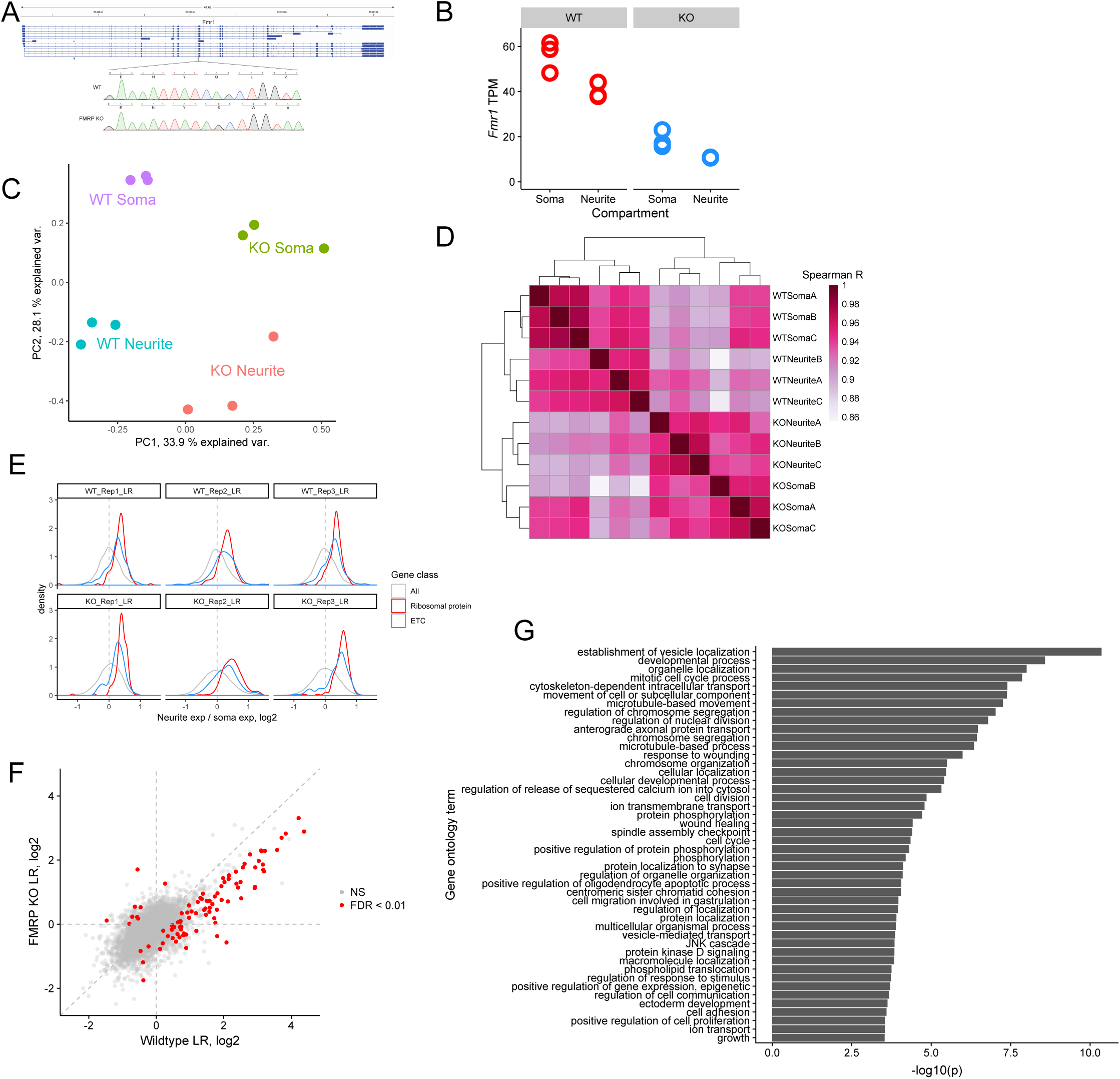
Analysis of soma/neurite fractionation sequencing in wildtype and FMRP null CAD cells. (A) Location of guide RNA directed against *Fmr1*. Sanger sequencing tracks of wildtype and knockout CAD cells are shown. In all alleles of the knockout cells, a single basepair deletion was created, leading to the creation of a premature stop codon two codons later. (B) RNA expression levels of *Fmr1* in the wildtype and FMRP null cells. (C) PCA analysis of gene expression values from the soma and neurite compartments of wildtype and FMRP null cells. (D) Correlation and clustering analysis of gene expression values from the soma and neurite compartments of wildtype and FMRP null cells. (E) LR values for all genes (gray), ribosomal protein genes (red), and genes that are part of the electron transport chain (blue). We have observed that RNAs encoding ribosomal protein genes and components of the electron transport chain are consistently neurite-enriched and therefore use them as markers to assess whether the soma/neurite fractionation was successful. (F) LR values for genes in wildtype and FMRP null cells. Those that met an FDR cutoff of 0.01, as determined by Xtail, are shown in red. (G) Gene ontology enrichments for genes with reduced neurite localization in FMRP knockout CAD cells.

**Figure S2.**
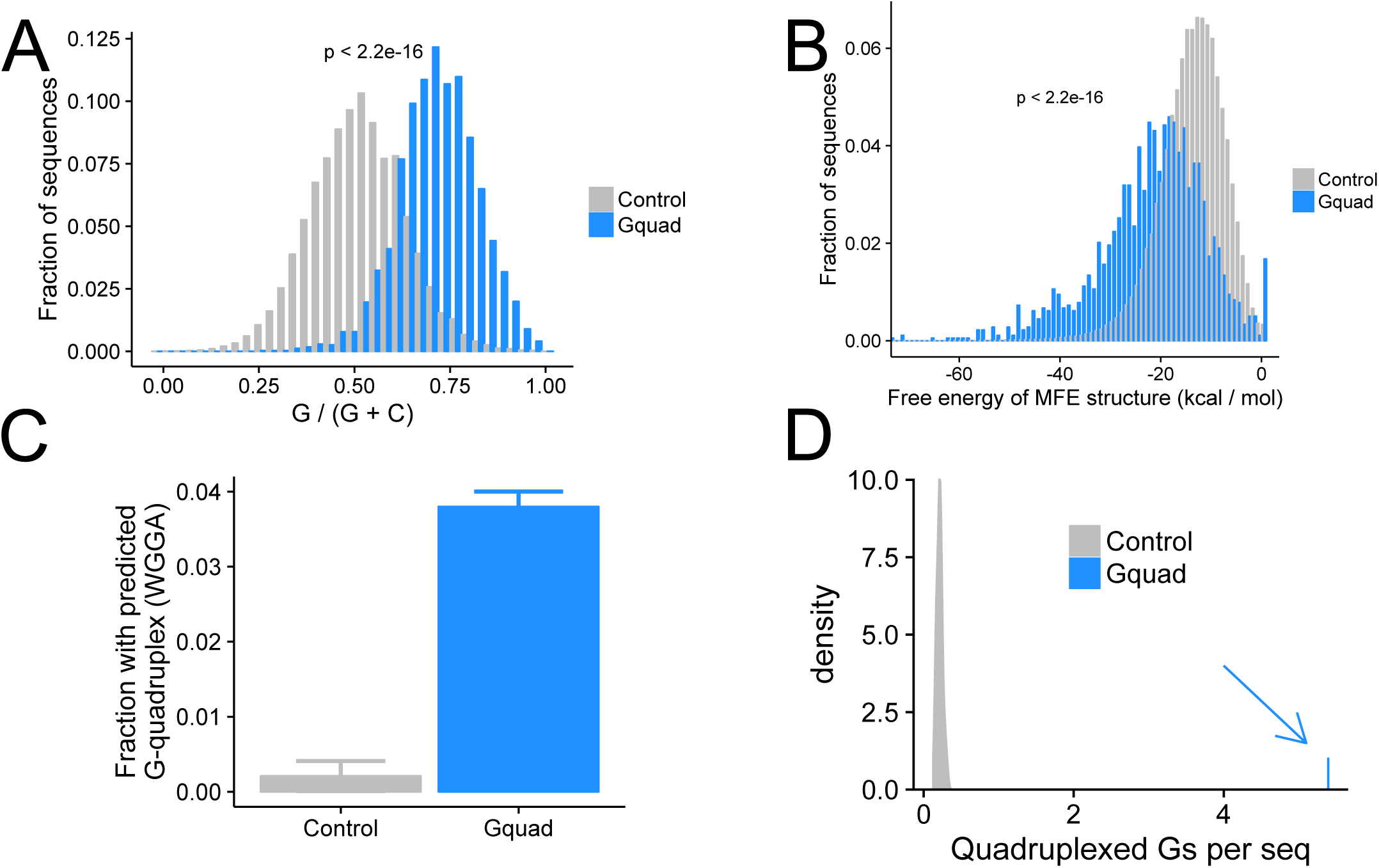
Analysis of experimentally defined G-quadruplex sequences. (A) Experimentally-defined G-quadruplex sequences contain more guanosine residues than control sequences drawn from the same genes. (B) Minimum free energy structures calculated using RNAfold show the G-quadruplex structures are significantly more stable than control sequences drawn from the same genes. (C) Experimentally defined G-quadruplex sequences contain many more regular expression-defined G-quadruplex sequences than controls. (D) Experimentally defined G-quadruplex sequences contain many more RNAfold-defined quadruplexed guanosine residues than control sequences.

**Figure S3.**
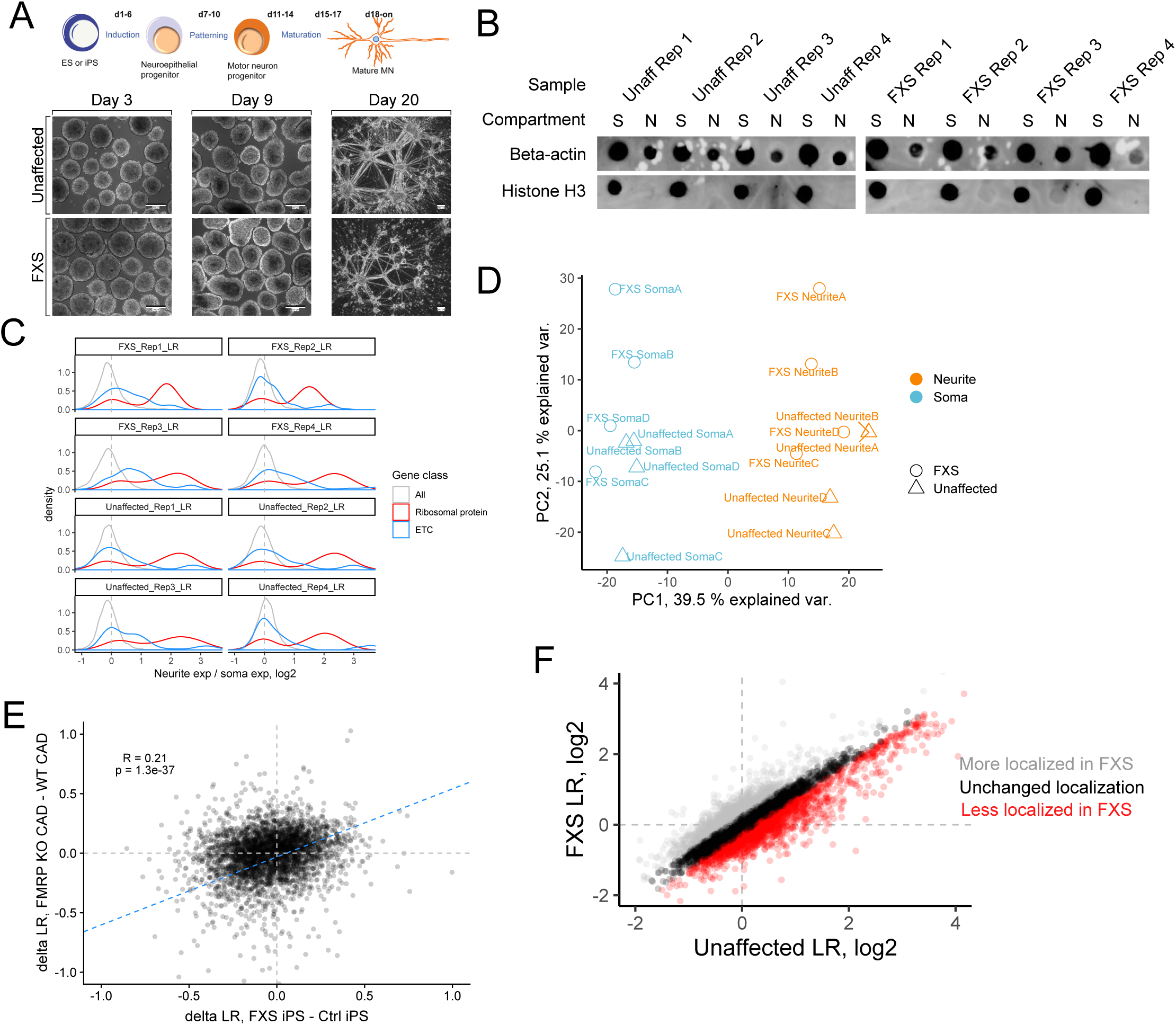
Fractionation of motor neurons differentiated from iPS cells derived from unaffected and FXS patients. (A) Schematic overview of motor neuron differentiation (top) and representative cell images (bottom). (B) Dot blot assaying the efficiency of motor neuron fractionations. Beta-actin is a marker of both soma (S) and neurite (N) fractions while Histone H3, being restricted to the nucleus, is a marker of soma fractions. (C) LR values for all genes (gray), ribosomal protein genes (red), and genes that are part of the electron transport chain (blue). (D) PCA analysis of gene expression values from the soma and neurite compartments of wildtype and FXS motor neurons. (E) Correlation between FMRP-dependent changes in LR values observed in mouse (FMRP-null CAD - wildtype CAD) and human (FXS motor neurons - unaffected motor neurons) neuronal cells. (F) LR values for genes observed in unaffected and FXS motor neurons. Genes whose log2 LR value changed by more than 0.25 were designated as either more localized in FXS (gray) or less localized in FXS (red).

**Figure S4.**
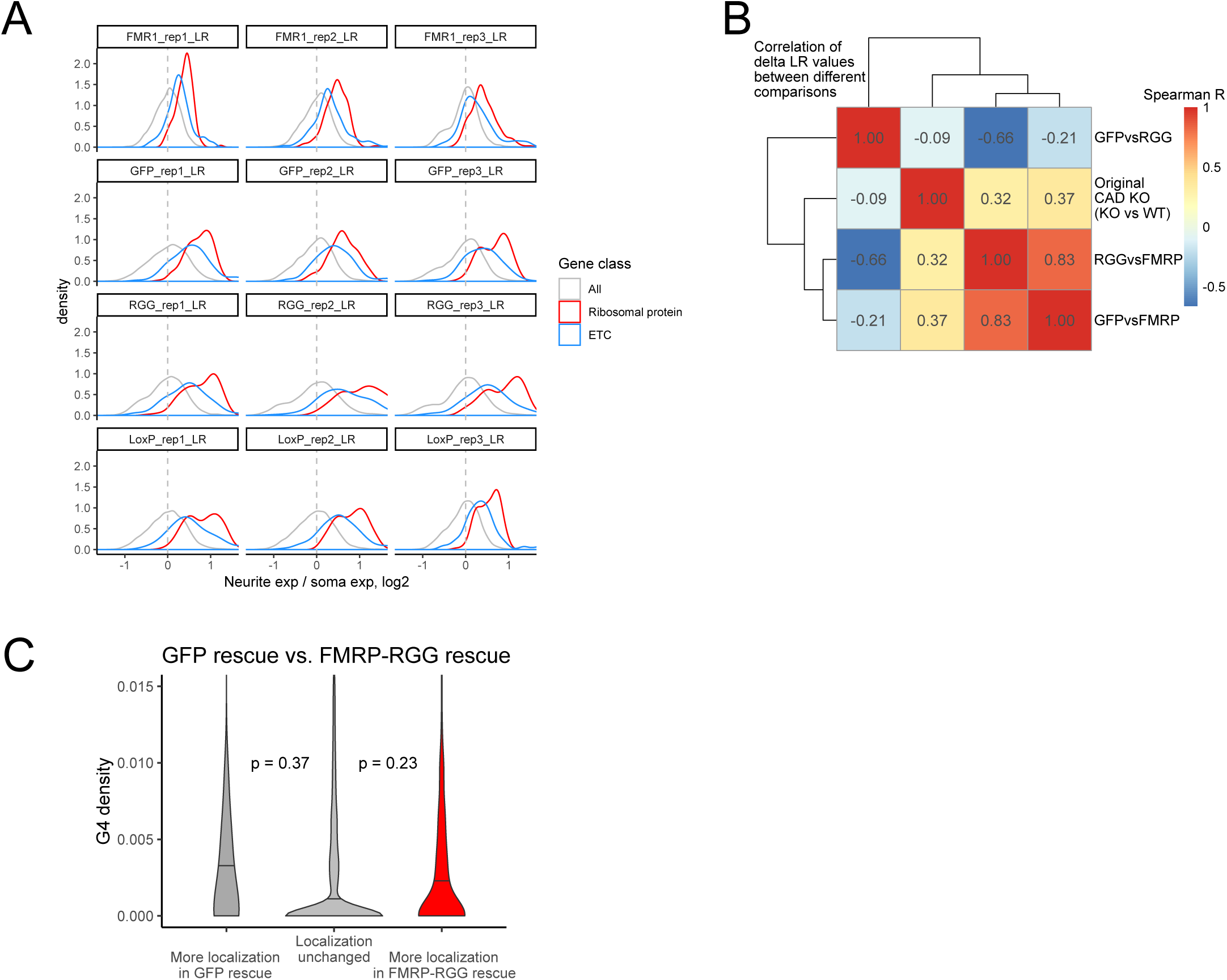
The RGG domain of FMRP is required for its efficient transport of G-quadruplex-containing transcripts. (A) LR values for all genes (gray), ribosomal protein genes (red), and genes that are part of the electron transport chain (blue). (B) Correlation of delta LR values (nonfunctional FMRP condition [GFP, RGG, or KO] - functional FMRP condition [WT or FMRP]) between experiments. (C) G-quadruplex density, as measured by RNAfold, in the 3′ UTRs of transcripts that were more localized in the GFP rescue cells (dark gray, left) or ΔRGG rescue cells (red, right). P values were calculated using Wilcoxon rank-sum tests. As in figure 2D, values represent the number of guanosine residues participating in quadruplex sequences per nt.

**Figure S5.**
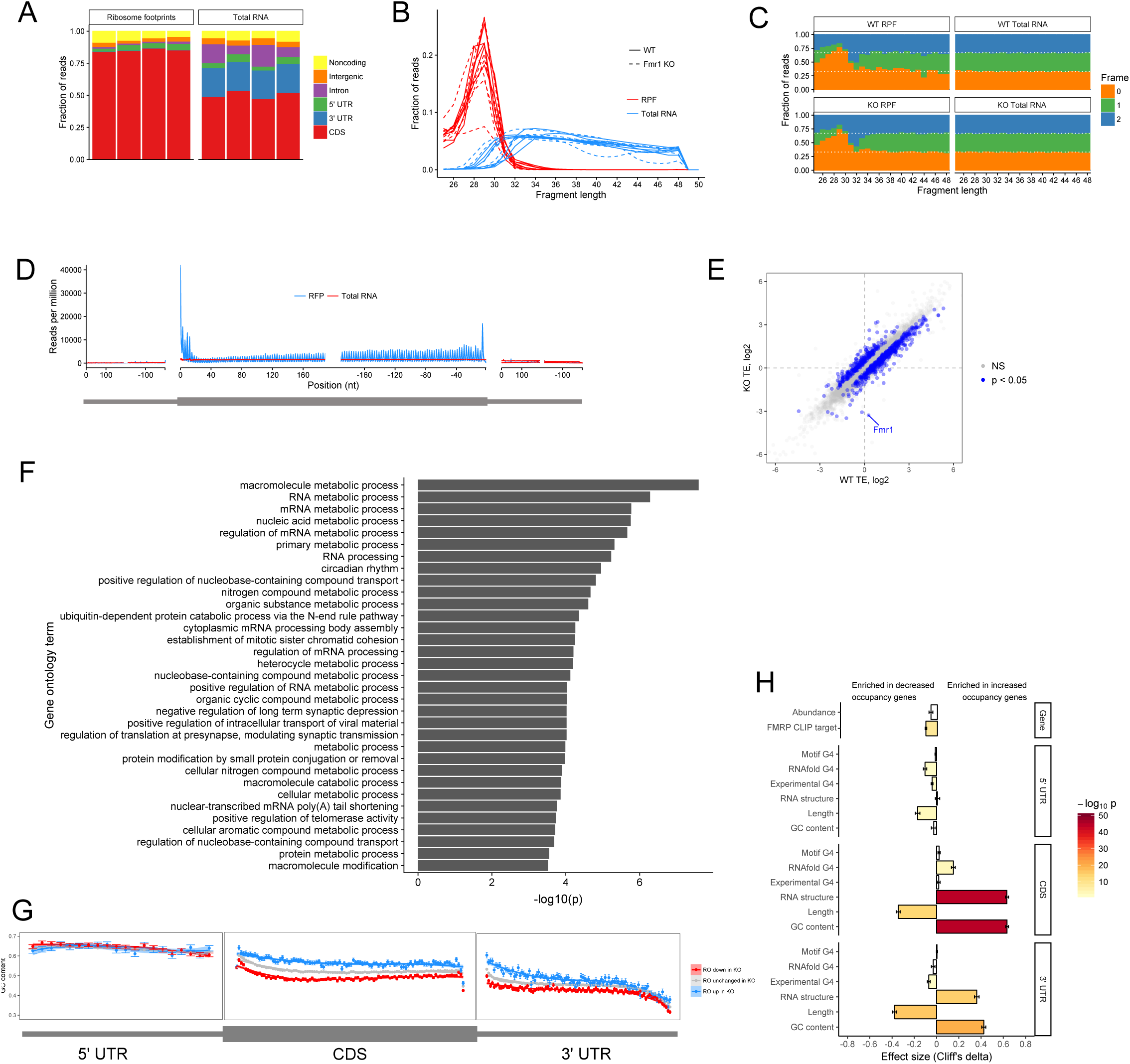
Ribosome profiling in wildtype and FMRP null CAD cells. (A) Fraction of reads that map to the indicated genomic regions. (B) Fraction of reads of that are the indicated lengths. (C) Fraction of reads that map to the indicated reading frames. (D) Metagene analysis of read densities across transcripts. Reads from ribosome protected fragments display a characteristic 3 nucleotide periodicity. (E) Ribosome occupancy values, as calculated by Xtail, in wildtype and FMRP null CAD cells. (F) Gene ontology analysis of the translational regulatory targets of FMRP in CAD cells. (G) Metagene analysis of GC content across transcripts whose ribosome occupancy increases (blue) or decreases (red) in FMRP null CAD cells compared to wildtype cells. (H) Effect sizes and significance of the relationships between transcript features and whether or not a transcript’s ribosome occupancy was increased or decreased by FMRP.

## SUPPLEMENTARY TABLES

**Table S1.** Xtail output for the differential localization of transcripts in wildtype and FMRP null CAD cells. All log2 fold change values are knockout / wildtype.

**Table S2.** Xtail output for the differential localization of transcripts in unaffected and FXS motor neurons. All log2 fold change values are FXS / unaffected.

**Table S3.** Xtail output for the differential localization of transcripts in FMRP null CAD cells rescued with either GFP or full length FMRP.

**Table S4.** Xtail output for the differential localization of transcripts in FMRP null CAD cells rescued with either ΔRGG or full length FMRP.

**Table S5.** Xtail output for the differential localization of transcripts in FMRP null CAD cells rescued with either ΔRGG or GFP.

**Table S6.** Xtail output for the differential ribosome occupancy of genes in wildtype and FMRP null CAD cells.

